# Variation at the common polysaccharide antigen locus drives lipopolysaccharide diversity within the *P. syringae* species complex

**DOI:** 10.1101/2020.03.31.019141

**Authors:** Jay Jayaraman, William T. Jones, Dawn Harvey, Lauren M. Hemara, Honour C. McCann, Minsoo Yoon, Suzanne L. Warring, Peter C. Fineran, Carl H. Mesarich, Matthew D. Templeton

**Affiliations:** Bioprotection Technologies, The New Zealand Institute for Plant and Food Research Limited, Auckland, New Zealand; Bioprotection Centre for Research Excellence, New Zealand; Bioprotection Technologies, The New Zealand Institute for Plant and Food Research Limited, Palmerston North, New Zealand; School of Biological Sciences, University of Auckland, New Zealand; Institute of Advanced Studies, Massey University, Auckland, New Zealand; Department of Microbiology and Immunology, University of Otago, Dunedin, New Zealand; School of Agriculture and Environment, Massey University, Palmerston North, New Zealand

**Author notes:** correspondence to Matthew Templeton. Max Planck Institute for Developmental Biology, Max-Planck-Ring 9, D-72076 Tübingen, Germany (Honour C. McCann).

## Abstract

The common polysaccharide antigen (CPA) from the lipopolysaccharide (LPS) component of cell walls from the species complex *Pseudomonas syringae* is highly variable both in structure and immunological specificity, but the genetic basis for this is not well understood. We have characterised the CPA locus from *P. syringae* pv. *actinidiae* (*Psa*). This locus has a modular structure with genes for both L- and D-rhamnose (Rha) biosynthesis and that of an unknown sugar. It also contains an operon coding for ABC transporter subunits, a bifunctional glycosyltransferase and an O-methyltransferase. This operon is predicted to have a role in *t*ransport, *e*longation and *t*ermination of the Rha backbone of the CPA oligosaccharide and is referred to as the TET operon. This is the first report of the identification of this operon in *P. syringae*. Two alleles of the TET operon were present amongst the different biovars of *Psa* and lineages of the closely related pathovar *P. syringae* pv. *actinidifoliorum*. This allelic variation was reflected in the electrophoretic properties of purified LPS from the different isolates. Gene knockout of the TET operon allele from biovar 1 and replacement with that from biovar 3, demonstrated the link between the genetic locus and the electrophoretic and immunogenic properties of the LPS molecules in *Psa*. Sequence analysis of the TET operon from a wide range of *P. syringae* and *P. viridiflava* isolates displayed a phylogenetic history which is incongruent with core gene phylogeny, but correlates with previously reported tailocin sensitivity, suggesting a functional relationship between LPS structure and tailocin susceptibility.

## INTRODUCTION

*Pseudomonas syringae* pv. *actinidiae* (*Psa*) is the causal agent of canker disease in kiwifruit (*Actinidia* Lindl spp.). Outbreaks of the disease were first observed in Japan and Korea in the 1980s and 1990s respectively, but isolates of the bacterium responsible for these outbreaks did not spread from their country of origin (Koh *et al.*, 1994; Serizawa *et al.*, 1989; Takikawa *et al.*, 1989). Between 2008 and 2010, a pandemic clone spread around the world, devastating the majority of regions growing cultivars of *A. chinensis* var. *chinensis* (gold kiwifruit) (Scortichini *et al.*, 2012). A large number of *Psa* isolates from a range of geographical origins have been sequenced, and phylogenies generated from the core genomes show that the three emergences of the disease are closely related, but form distinct clades (McCann *et al.*, 2017; McCann *et al.*, 2013). Single Nucleotide Polymorphism (SNP) analysis indicates that the recent pandemic isolate originated in China, but the location of the source population of this pathovar has not been identified (McCann *et al.*, 2017). Despite the close phylogenetic relationship between *Psa* isolates, their accessory genomes vary significantly in their effector complement and secondary metabolite portfolios (Butler *et al.*, 2013; Marcelletti *et al.*, 2011; Mazzaglia *et al.*, 2012; McCann *et al.*, 2013). Based on both the phylogeny of the core genome and accessory gene variation, *Psa* isolates have been designated as biovars (BVs) (Cunty *et al.*, 2015; Vanneste *et al.*, 2013). The isolates that only cause leaf spots on kiwifruit, initially described as BV4, are closely related to *Psa*, but have now been given their own pathovar designation; *P. syringae* pv. *actinidifoliorum* (*Pfm*), and consist of four distinct lineages (Cunty *et al.*, 2015). Recently, two more *Psa* BVs have been discovered in Japan (Fujikawa and Sawada, 2016; Sawada *et al.*, 2016). Variation in the accessory genome of *P. syringae* pathovars and *Psa* BVs, and its role in host specificity has been well documented (Dillon *et al.*, 2019a; McCann *et al.*, 2013). In addition, *P. syringae* isolates have been shown to have a high degree of structural and serological variation in their lipopolysaccharide (LPS) (Zdorovenko and Zdorovenko, 2010). However, the genetic basis for this is not well understood. Both bacteriophages and tailocins utilise LPS as a receptor to recognise and bind to their host. Tailocins are derivatives of bacteriophages, comprising predominantly the tail proteins that function to depolarise the bacterial membrane (Riley and Wertz, 2002). Isolates of *P. syringae* use tailocins (also known as R-type syringacins) to target and outcompete closely related strains that presumably occupy a similar ecological niche (Hockett *et al.*, 2015). It has been shown recently that there is a high degree of variation in the sensitivity to tailocins within *P. syringae* (Baltrus *et al*., 2019). Understanding the molecular basis of this variation is important if we are to implement novel methods such as the use of phage therapy or tailocins for biological control of this pathogen (Baltrus *et al.*, 2019; Frampton *et al.*, 2015; Frampton *et al.*, 2014; Pinheiro *et al.*, 2019; Rooney *et al.*, 2019).

LPS are complex glycolipids that make up the surface leaflet of the outer membrane of gram-negative bacteria, forming a physical protective barrier. LPS have three distinct domains: Lipid A that tethers the molecule to the outer membrane, the core domain, and finally the O-polysaccharide (OPS). The last domain includes the O-specific antigen (OSA) and/or the common O-polysaccharide antigen (CPA), and is structurally the most variable component of the LPS. The distinction between the two OPS polysaccharides is that the CPA is exported via an ABC transporter-dependent pathway, while OSA synthesis follows the Wzx/Wzy-dependent pathway (Raetz and Whitfield, 2002). In many bacterial species, the OPS are responsible for the immunological variation between isolates and, for human and animal pathogens, subspecies classification is based on serotypes (DebRoy *et al.*, 2016; King *et al.*, 2009). Similarly, the structures of OPS in plant-pathogenic bacteria such as *P. syringae* are highly variable at the pathovar level (Ovod *et al.*, 1997a). For *P. syringae*, the OPS appears to be comprised of the CPA (Kutschera *et al.*, 2019; Mesarich *et al.*, 2017) with a backbone consisting of a tri- or tetra-saccharide repeating unit, comprising either L- or D-Rhamnose (Rha), or a combination of the two, with various linkages. These are decorated with side chains consisting of different sugars such as *N*-acetyl-glucosamine (GlcNAc), fucose (Fuc), Rha, and *N*- acetyl-fucosamine (FucNAc) (Ovod *et al.*, 1997a; Zdorovenko and Zdorovenko, 2010). The OPS can also be methylated or acetylated to varying degrees (Zdorovenko *et al.*, 2001). At least nine serotypes of *P. syringae* have been identified, and each differs in the structure of the OPS backbone and side-chain modifications (Zdorovenko and Zdorovenko, 2010). Although considerable effort in the past was devoted to attempts at relating serotype to pathovar identification and taxonomy, correlations were not consistently observed (Ovod *et al.*, 1997b). Little is known about the biosynthesis of LPS in *P. syringae* or the genetic basis for the observed structural and immunological variability within the genus.

Transposon mutagenesis of *Psa* BV3 using Tn*5* identified a set of mutations with a rough (R) colony phenotype (Mesarich *et al.*, 2017). This phenotype is often associated with mutations in LPS biosynthesis. Indeed, the majority of the Tn*5* inserts in these R-LPS mutants mapped to a gene cluster in *Psa* orthologous to the CPA biosynthetic pathway from *P. aeruginosa* (Mesarich *et al.*, 2017). Furthermore, LPS was not observed in the mutants, suggesting CPA is the sole OPS in *Psa* (Hockett *et al.*, 2017). Other recent papers have directly identified components of CPA biosynthesis in *P. syringae* through gene knockouts (Kutschera *et al.*, 2019) and increased tailocin resistance (Kandel *et al.*, 2019; Kutschera *et al.*, 2019).

Here, we investigated the genetics of LPS biosynthesis in *Psa*. We characterised the CPA locus and the orthologous region in the different *Psa* BVs, *P. syringae* and the closely related *P. viridiflava* isolates. We show an operon from this locus is hyper-variable and accounts for the high degree of structural and immunological variation observed in this species complex. Furthermore, the relationship between this component of the CPA locus and sensitivity to tailocins was highly correlated.

## RESULTS

### Major LPS biosynthesis genes in *Psa* BV3 are shared with *P. aeruginosa*

In *P. aeruginosa* PAO1, the genes and operons responsible for the biosynthesis of Lipid A, the core LPS and the OPS tend to be co-located and have been well annotated and characterised (King *et al.*, 2009; Lam *et al.*, 2011). We identified the key loci containing the majority of the LPS biosynthetic genes in *Psa* BV3 using BLASTx with *P. aeruginosa* PAO1 orthologs (Figure 1A & Table 1). Amino acid similarity to *P. aeruginosa* PAO1 orthologs of the Lipid A, core LPS, and Rha biosynthetic pathways in the CPA locus is relatively high (Table 1). In contrast, *Psa* genes putatively coding for other components of the CPA, such as the *wzm*/*wzt*/*wbpX* orthologues, are more divergent and have poor or no significant BLASTx hits. Evidence for an OSA biosynthetic cluster was not found, apart from genes in the *wbpK*-*M* locus, which is also involved in CPA biosynthesis (Table 1). The co-location of genes annotated as ABC transporters and glycosyltransferases (GT) with both the D- and L-Rha biosynthetic pathways suggests that the CPA is the predominant OPS in *Psa* BV3 (Figure 1A).

**Table 1.**
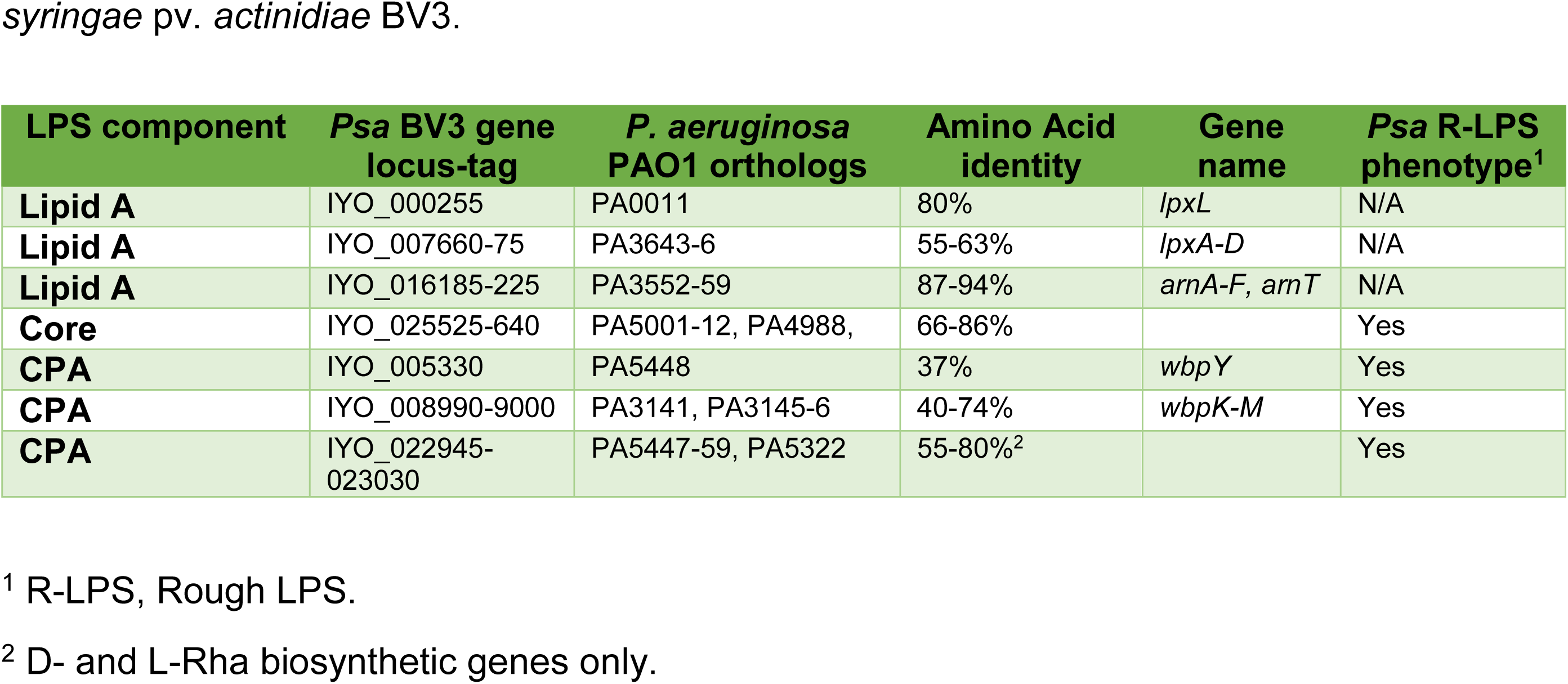
Loci involved in lipopolysaccharide (LPS) biosynthesis in *Pseudomonas syringae* pv. *actinidiae* BV3.

| LPS component | Psa BV3 gene locus-tag | *P. aeruginosa* PAO1 orthologs | Amino Acid identity | Gene name | Psa R-LPS phenotype<sup>1</sup> |
| --- | --- | --- | --- | --- | --- |
| **Lipid A** | IYO_000255 | PA0011 | 80% | *lpxL* | N/A |
| **Lipid A** | IYO_007660-75 | PA3643-6 | 55-63% | *lpxA-D* | N/A |
| **Lipid A** | IYO_016185-225 | PA3552-59 | 87-94% | *arnA-F, arnT* | N/A |
| **Core** | IYO_025525-640 | PA5001-12, PA4988, | 66-86% |  | Yes |
| **CPA** | IYO_005330 | PA5448 | 37% | *wbpY* | Yes |
| **CPA** | IYO_008990-9000 | PA3141, PA3145-6 | 40-74% | *wbpK-M* | Yes |
| **CPA** | IYO_022945-023030 | PA5447-59, PA5322 | 55-80%<sup>2</sup> |  | Yes |

<sup>1</sup> R-LPS, Rough LPS.
<sup>2</sup> D- and L-Rha biosynthetic genes only.

**Figure 1.**
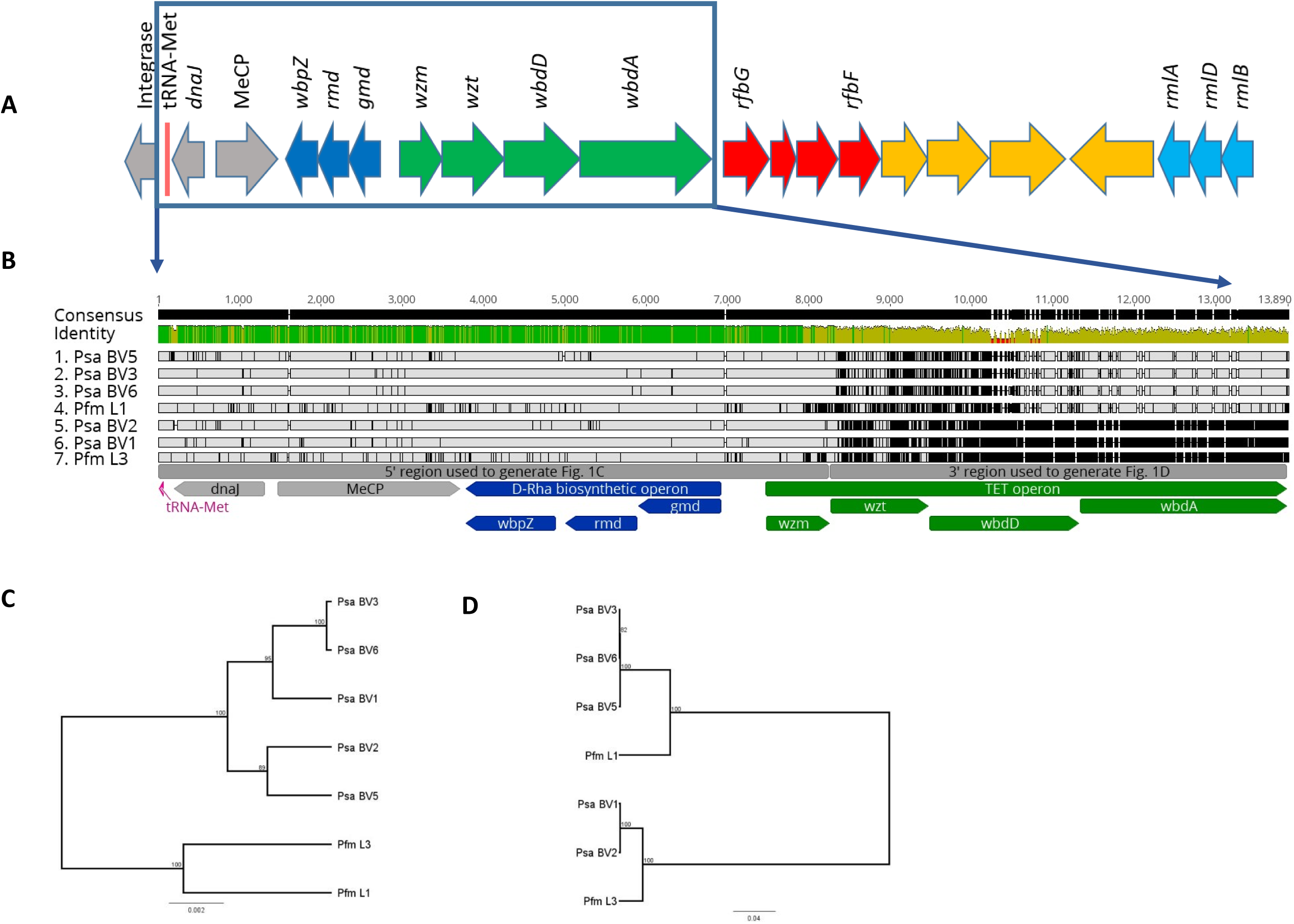
The common polysaccharide antigen (CPA) locus from *Pseudomonas syringae* pv. *actinidiae* (*Psa*) and *P. syringae* pv. *actinidifoliorum.* **(A)** The genetic structure of the CPA locus in *Psa* ICMP18884 (BV3). The four operons annotated are: the D-Rha biosynthesis operon (dark blue), CPA transport, extension, and termination (TET) operon (green), synthesis of unknown sugar operon (red), L-Rha synthesis operon (light blue), neighbouring syntenic genes that are not believed to be part of this functional cluster are shaded grey or yellow. **(B)** Comparative analysis of the conserved D-Rha biosynthesis and polymorphic TET operons of *Psa* and *Pfm*. The CPA locus beginning at the syntenic conserved tRNA-Met through *wbdA* of the TET locus for representatives of the *Psa* biovars (BV1, ICMP 9617; BV2, ICMP 19071; BV3, ICMP 18884; BV5, MAFF 212063; and BV6, MAFF 212141) and *Pfm* lineages (L1, ICMP 18803; L3, ICMP 18807) aligned in Geneious v10 with consensus identity displayed above the aligned sequences. Flanking DnaJ and MeCP genes are annotated in grey, D-Rha biosynthesis operon and its genes are annotated in dark blue, and TET operon and its genes annotated in green. **(C)** UPGMA tree of the CPA locus from 5’ region up to and including *wzm* in (A). **(D)** UPGMA tree of the CPA locus from 3’ region from *wzt* to *wbdA* in (B).

### Bioinformatic identification of the CPA locus in *Psa* BV3

Identifying the CPA locus in *P. syringae* has been challenging due to the lack of similarity to *wzm*/*wzt*/*wbpX* orthologues (Kutschera *et al.*, 2019). Previous work identified five R-LPS mutants by Tn*5* mutagenesis that had inserts in a region with homology to the CPA biosynthetic pathway from *P. aeruginosa* PAO1 (Mesarich *et al.*, 2017). Bioinformatic analysis of this region revealed a locus of 18 genes (Figure 1A). These genes were arranged in operons predicted to be involved in the synthesis of L-Rha, D-Rha, an unknown sugar, an ABC transporter complex involved in the transport of the CPA, a bifunctional glycosyltransferase, an O-methyltransferase, and a region of unpredicted function (Table 2). Of the two genes annotated as ABC transporters, the first (IYO_023015) has homology to the ABC permease superfamily gene *wzm*, while the second (IYO_023010) has an N-terminal ATPase domain with a C-terminal *wzt*-like domain (hereafter called *wzt*). The latter domain binds to the non-reducing terminal modification of the sugar chain and is thus specific for the transported molecule (Cuthbertson *et al.*, 2007). The GT gene (IYO_023000) on the same operon as the two ABC transporters has two GT4 domains, implying it catalyses two distinct glycosyl transfer reactions. It is homologous to *wbdA* (formerly known as *mtfB*), a functionally characterised mannosyltransferase in *E. coli*, which has been shown to direct the extension of the O9-specific polysaccharide chain by synthesising, then polymerising, the tetra-saccharide repeat (Liston *et al.*, 2015). The adjacent gene (IYO_023005) on the *Psa* BV3 operon is annotated as an O-methyltransferase. In *E. coli*, the LPS oligosaccharide chain extension, catalysed by WbdA, is terminated by a dual methyltransferase/kinase enzyme (WbdD) (Mann *et al.*, 2019). It appears therefore that this operon possesses genes required for the *<u>t</u>*ransport, *<u>e</u>*xtension, and *<u>t</u>*ermination of the backbone oligosaccharide chain of the CPA in *Psa* BV3 (hereafter referred to as the TET operon). This is the first time this operon has been identified in *P. syringae*.

**Table 2.** Gene annotations of *Pseudomonas syringae* pv. *actinidiae* BV3 common LPS locus.

| Gene annotation | *P. aeruginosa* | *E. coli* | Function | Gene ID | Protein_ID | R-LPS<sup>1</sup> |
| --- | --- | --- | --- | --- | --- | --- |
| dTDP-glucose 4,6-dehydratase |  | *rmlB* | L-Rha biosynthesis | IYO_022945 | AKT32330.1 |  |
| dTDP-4-dehydrorhamnose reductase |  | *rmlD* | L-Rha biosynthesis | IYO_022950 | AKT32331.1 |  |
| glucose-1-phosphate thymidyltransferase |  | *rmlA* | L-Rha biosynthesis | IYO_022955 | AKT32332.1 |  |
| hypothetical protein |  |  | unknown | IYO_022960 | AKT32333.1 |  |
| hypothetical protein |  |  | unknown | IYO_022965 | AKT32334.1 |  |
| amine oxidase |  |  | unknown | IYO_022970 | AKT32335.1 |  |
| membrane protein |  |  | unknown | IYO_022975 | AKT32336.1 |  |
| glucose-1-phosphate cytidyltransferase |  | *rfbF* | synthesis of unknown sugar | IYO_022980 | AKT32337.1 |  |
| glycosyl transferase |  |  | synthesis of unknown sugar | IYO_022985 | AKT32338.1 |  |
| NAD-dependent epimerase/dehydratase |  | *galE* | synthesis of unknown sugar | IYO_022990 | AKT32339.1 |  |
| CDP-glucose 4,6-dehydratase |  | *rfbG* | synthesis of unknown sugar | IYO_022995 | AKT32340.1 |  |
| glycosyl transferase family 1 | *wbpX* | *wbdA* | chain elongation | IYO_023000 | AKT32341.1 |  |
| SAM-dependent methyltransferase |  | *wbdD* | chain termination | IYO_023005 | AKT32342.1 | Yes |
| sugar ABC transporter ATP-binding protein | *wzt* | *wzt* | transport | IYO_023010 | AKT32343.1 | Yes |
| ABC transporter | *wzm* | *wzm* | transport | IYO_023015 | AKT32344.1 | Yes |
| **GDP-D-mannose dehydratase** | *gmd* | *gmd* | D-Rha biosynthesis | IYO_023020 | AKT32345.1 | Yes |
| **GDP-6-deoxy-D-lyxo-4-hexulose reductase** | *rmd* | *rmd* | D-Rha biosynthesis | IYO_023025 | AKT32346.1 |  |
| **glycosyl transferase family 1** | *wbpZ* |  | D-Rha biosynthesis | IYO_023030 | AKT32347.1 | Yes |

<sup>1</sup> R-LPS, Rough LPS.

### The TET operon from *Psa* and *Pfm* is encoded by two different alleles

The CPA locus from representatives of all five *Psa* BVs and two lineages of *Pfm* were identified and compared. The TET operon, genes for D-Rha biosynthesis, and regions coding for a chemotactic receptor, a DnaJ molecular chaperone and a tRNA-Met were shared and could be aligned (Figure 1B). The latter three genes were syntenic in all genomes but are not part of the CPA locus. The region including the D-Rha biosynthetic pathway and the first gene (*wzm*) in the TET operon (5’ region) is well conserved with few SNPs and a phylogenetic tree (Figure 1C) closely matched the topology of a tree generated from seven conserved genes (Sawada *et al.*, 2016). In contrast, the 3’ region, from the gene encoding the ABC transporter ATPase (*wzt*) of the TET operon is more variable, representing two groups of divergent sequences. One group includes *Psa* BVs 1 and 2, and *Pfm* L3, while the other includes *Psa* BVs 3, 5, and 6, and *Pfm* L1. A phylogenetic tree of this region (Figure 1D) resolved into two clades that did not match the topology of Figure 1C, or the phylogeny generated from the core genome (Supplementary Figure S1). These results suggest that variation at the TET operon is driving diversity at the CPA locus in *Psa* and *Pfm*.

### LPS from *Psa* and *Pfm* has different biochemical properties

To assess whether genetic variation in the TET operon results in variation of the biochemical properties of LPS, we extracted LPS from *Psa* and *Pfm* using the method of Westphal and Jann (1965). Resolution of the extracted LPS by SDS-PAGE revealed the classic ladder pattern characteristic of bacterial LPS molecules, a reflection of the length and charge micro-heterogeneity of the O-linked saccharides (Jann *et al.*, 1975). Comparison of the LPS between the five different BVs of *Psa* and two lineages of *Pfm* revealed two patterns based on differences in the molecular weight range and spacing between bands (Figure 2A). LPS from *Psa* BV1, BV2 and *Pfm* L3 formed one group, which was characterised by a broad range of molecular weights, especially in the higher molecular mass range. In contrast, *Psa* BV3, BV5, BV6 and *Pfm* L1 formed a second group that had a lower range of molecular weights and narrower spacing between bands (Figure 2A). The two electrophoretic pattern groups reflect the broad clusters evident in the TET operon tree (Figure 1D), suggesting the TET operon plays a significant role in defining the electrophoretic properties of LPS. Banding patterns within a group were not identical, suggesting that they may differ in decoration of the back-bone with side-chain sugars and/or acylation. Therefore, the genetic and biochemical data indicate that the CPA locus identified in *Psa* and *Pfm* is likely responsible for LPS biosynthesis in these *P. syringae* pathovars.

**Figure 2.**
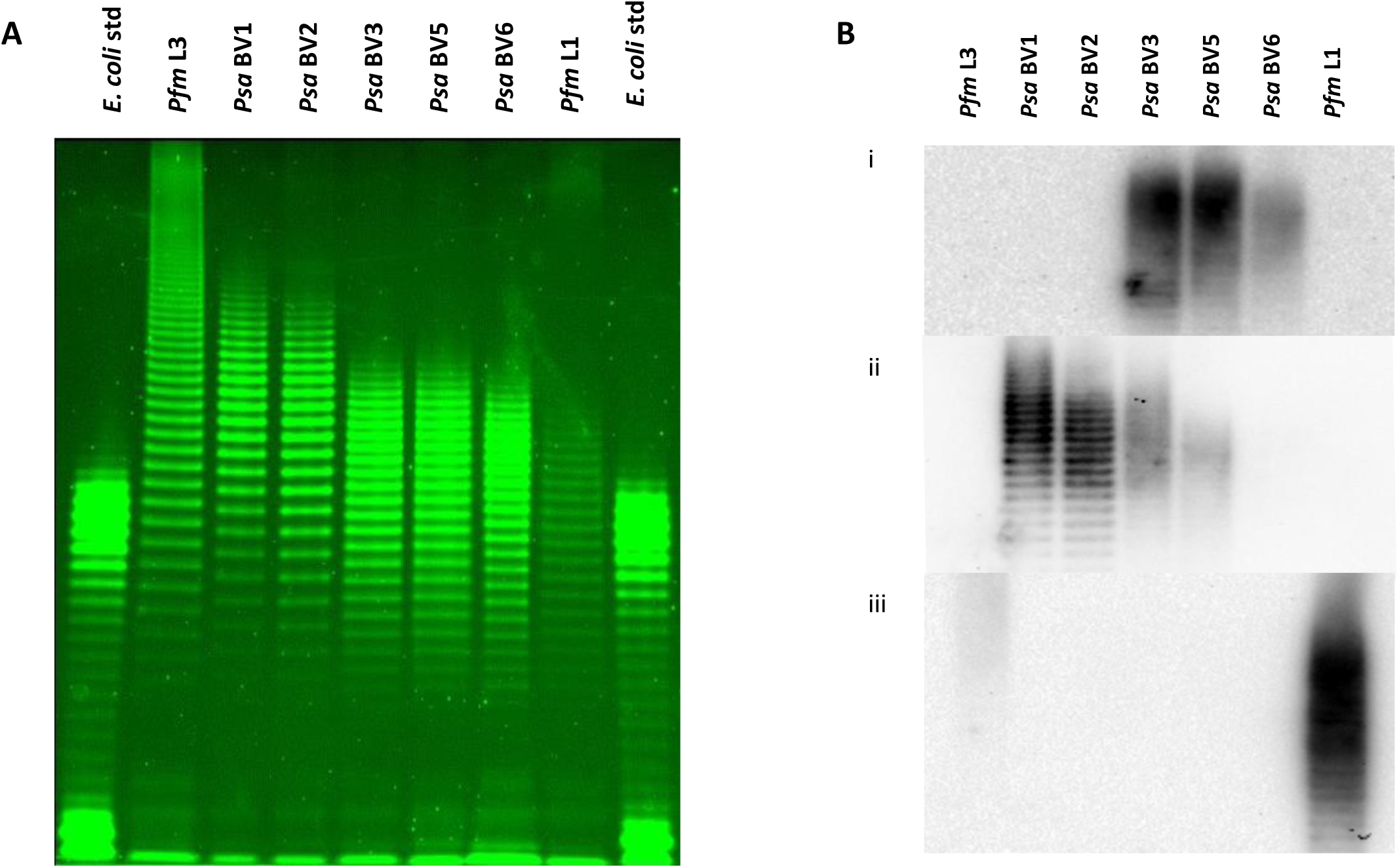
LPS profiles of *Pseudomonas syringae* pv. *actinidiae* (*Psa*) and *P. syringae* pv. *actinidifoliorum* (*Pfm*). **(A)** Pro-Q™ Emerald 300 staining of lipopolysaccharide (LPS) preparations on a NuPAGE 4–12% gradient bis-tricine SDS gel from representatives of *Pfm* lineages (L3 ICMP 18807, lane 2; L1 ICMP 18803, lane 8), *Psa* biovars (BV1 ICMP 9617, lane 3; BV2 ICMP 19071, lane 4; BV3 ICMP 18884, lane 5; BV5 MAFF212063, lane 6; BV6 MAFF 212141, lane 7), with *Escherichia coli* standards (Std; lanes 1 and 9). A total of 0.5–15 μL LPS extract for each sample (normalized by trial runs) was applied per lane. **(B)** Western blot of LPS extracts from representatives of *Psa* biovars and *Pfm* lineages run on a NuPAGE 4– 12% gradient bis-tricine SDS gel, obtained by probing with polyclonal antibodies to *Psa* BV3 ICMP 18884 **(Bi)**, *Psa* BV1 ICMP 9617 **(Bii)**, or *Pfm* L1 ICMP 18803 **(Biii)** in a 5000:1 ratio.

### LPS from *Psa* and *Pfm* are immunologically distinct

Western blots of the various LPS preparations for *Psa* and *Pfm* isolates were probed with each isolate-specific polyclonal antibody preparation (Figure 2B). All purified polyclonal antibodies showed a high degree of specificity, with little cross-reactivity against closely related isolates. The anti-*Psa* BV3 antibody preparation was the most specific, reacting with BV3, BV5, and only weakly to BV6 LPS (Figure 2Bi). The weak binding to the *Psa* BV6 LPS and the slightly different *Psa* BV6 LPS profile (Figure 2A) suggest a subtle difference in the CPA decoration. There was no cross-reactivity of the *Psa* BV3 antibody to the *Psa* BV1, BV2, or *Pfm* LPS preparations (Figure 2B). The lack of binding to *Pfm* L1 LPS (which appears more similar to BV3 LPS than *Pfm* L3) is likely due to the fact that antibodies that bind to both BV3 and *Pfm*L1 were depleted by the affinity chromatography purification process used. The anti-BV1 antibody preparation reacted strongly with both *Psa* BV1 and BV2 LPS, as expected based on the close genetic identity between their TET loci (Figure 2Bii). There was also weak cross-reactivity of *Psa* BV1 with *Psa* BV3 and BV5 LPS preparations (Figure 2Bii). Interestingly, the *Pfm* L1 antibody had a slight cross-reactivity with *Pfm* L3, but not with the *Psa* isolates in the same grouping (Figure 2Biii). These results further support the different grouping of LPS between these strains, but also indicate that antibody specificity is influenced by both backbone structure and side-chain decorations.

### The TET operon determines the variability of LPS banding in *Psa*

To confirm whether the TET operon, expressing the ABC transporter (*wzt*), putative oligosaccharide chain elongation GT (*wbdA*), and the O-methyl transferase (*wbdD*), are responsible for the different LPS profiles, this operon was deleted in *Psa* BV1 (Figure 3A; dark green region, Supplementary Figure S2) and replaced with that from *Psa* BV3 (Figure 3A, blue region, Supplementary Figure S2). The *Psa* BV1 TET operon knock-out (Δ*LPS* KO) lacked LPS and possessed a rapid sedimentation phenotype, but had a very weak R-LPS colony morphology, as observed for the BV3 mutants (Figure 3B; Supplementary Figures. S3A & S4). *Psa* BV1 with the BV3 LPS knocked back in (four knock-in [KI] lines), but not for the KO revertants that failed to retain this region, restored these LPS-associated phenotypes (Figure 3B, Supplementary Figures. S3B & S4). Furthermore, the LPS profile was identical to that of wild-type BV3, proving this locus codes for the genes involved in generating the BV-specific LPS oligosaccharide ladder. A Western blot of the LPS gel of these strains was probed with the *Psa* BV3 antibody (Figure 3C), which showed that transfer of the TET locus from *Psa* BV3 to the *Psa* BV1 Δ*LPS* KO also transferred the antibody specificity. In summary, this genetic TET-swap analysis provided direct evidence that these genes were necessary and sufficient for the differences in LPS between these strains.

**Figure 3.**
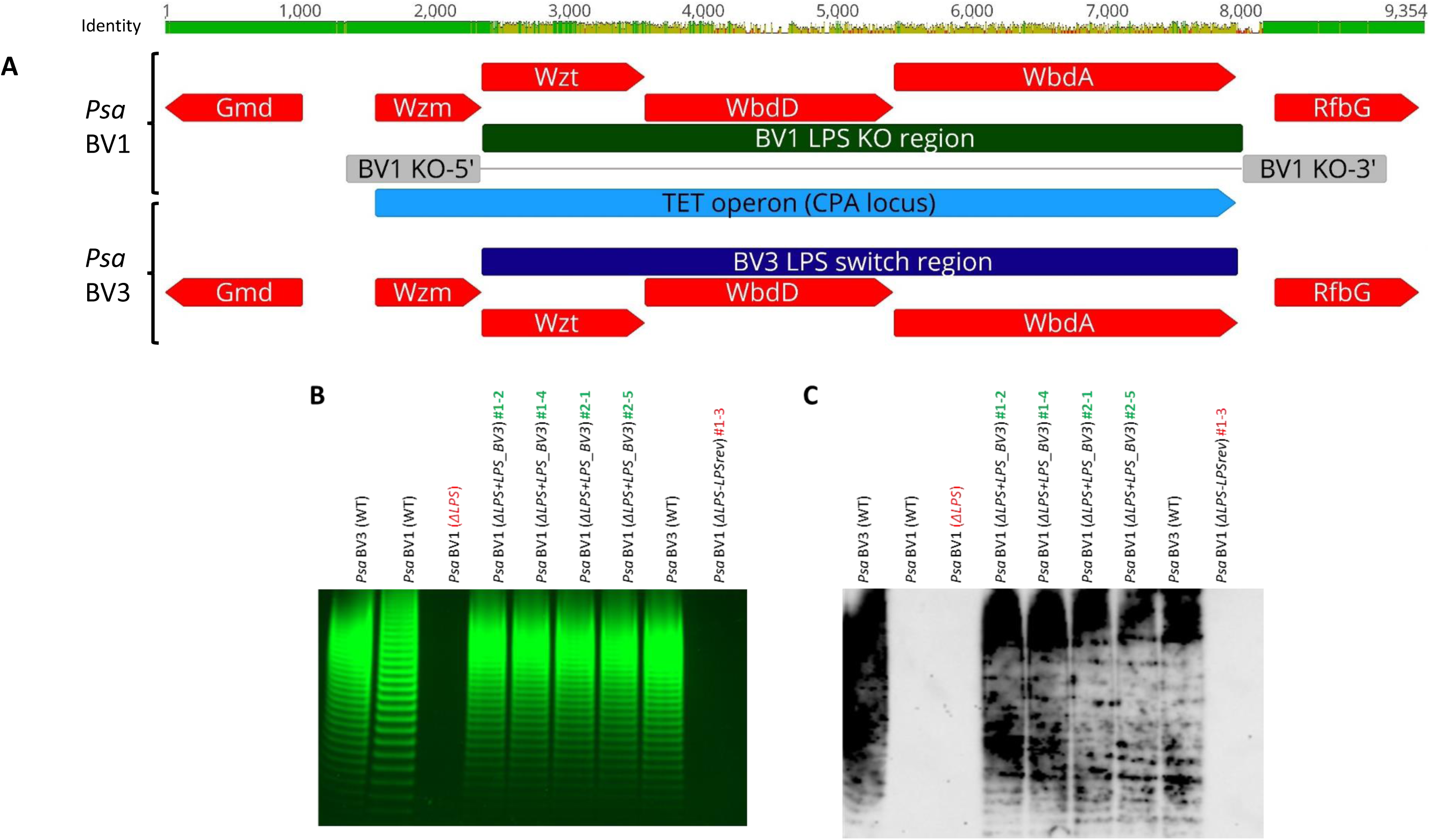
Effects of CPA TET operon swap on LPS banding and immunological recognition in *Pseudomonas syringae* pv. *actinidiae* (*Psa*). **(A)** Genetic polymorphism of the TET operon for *Psa* BV1 (ICMP 9617) versus BV3 (ICMP 18884). Genetic sequences aligned in Geneious v10. Genome sequences are indicated by light grey bars with associated translated regions as black bars, operons are annotated in light blue, genes in light green with gene names indicated, coding sequences in red with predicted biochemical function indicated, the original TET operon from BV1 indicated in dark green with upstream (KO-5’) and downstream (KO-3’) regions for generating the LPS TET operon knock-out in grey, and the TET operon swap region by knock-in from BV3 indicated in dark blue. **(B)** LPS profiles of *Psa* BV3 (lane 1 and lane 8), BV1 (lane 2), BV1 TET operon knockout (Δ*LPS*; lane 3), four BV1-to-BV3 TET operon complemented isolates (Δ*LPS*+*LPS*_BV3 1-2, 1-4, 2-1, and 2-5; lanes 4–7), and one BV1 revertant to knock-out isolates (ΔLPS-LPSrev 1-3; lane 9). Pro-Q™ Emerald 300 staining of LPS preparations on a 4–12% gradient Tris-glycine SDS gel. LPS extract (15 μL) for each sample was applied per lane. **(C)** Immuno-blot of LPS extracts from wild-type, knock-out (KO), KO-complemented, and KO-revertant strains run on a gradient SDS-PAGE gel as for (A), probed with polyclonal antibodies to *Psa* BV3 in a 5000:1 ratio.

### LPS is required for full pathogenicity of *Psa* in kiwifruit plantlets

Previously, LPS mutations have been found to affect host colonization by *P. syringae* (Kutschera *et al.*, 2019). To investigate the role of LPS in the virulence or pathogenicity of *Psa*, we infected kiwifruit plantlets with *Psa* BV3 R-LPS Tn*5* mutants (Δ*wbpL*, Δ*wzm*, Δ*wbdD*, Δ*wzt*, and Δ*gmd*) (Mesarich *et al.*, 2017) and the *Psa* BV1 knock-out (Δ*LPS*), two knock-in (Δ*LPS+LPS_BV3*) and two knock-out revertant (Δ*LPS-LPSrev*) mutants, along with their wild-type counterparts. All LPS mutants showed at least a log reduction in growth *in planta* compared to wild-type strains, indicating that LPS is required for host colonization (Figure 4A-B) and symptom development (Supplementary Figure S5A-B). Meanwhile, the BV1Δ*LPS+LPS_BV3* strains with complemented LPS production restored wild-type growth to the Δ*LPS* mutant (Figure 4B; Supplementary Figure S5B) and this did not occur in the strains that had reverted to Δ*LPS*. This correlated with the strains’ ability to stay suspended in inoculation buffer (Supplementary Figure S3A-B).

**Figure 4.**
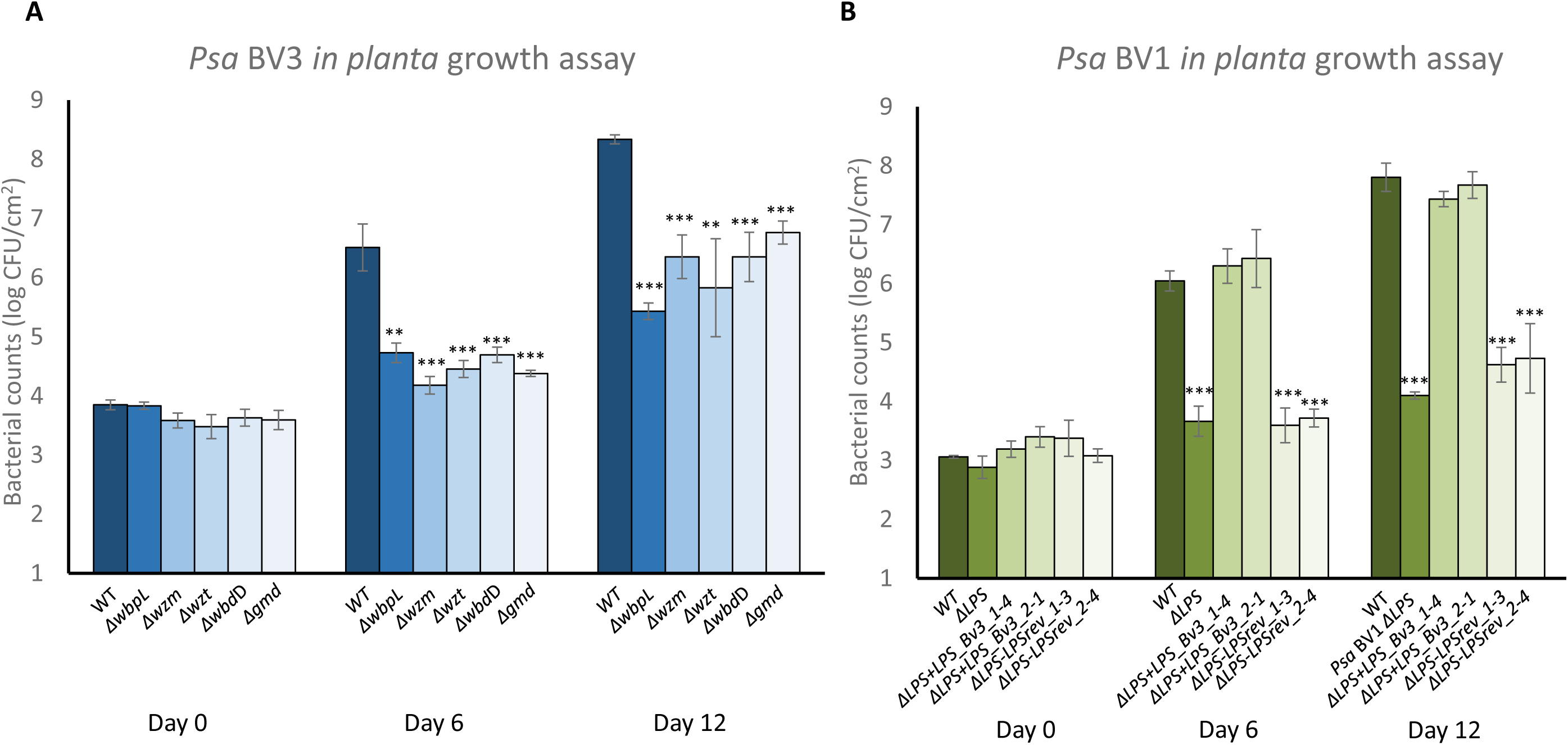
The CPA TET operon is required for full pathogenicity of *Pseudomonas syringae* pv. *actinidiae* (*Psa*) in host plants. **(A)** *P*sa BV3 or BV3 R-LPS mutants (Mesarich *et al.*, 2017) were flood-inoculated at ∼10^6^ CFU/mL on *Actinidia chinensis* var. *chinensis* ‘Zesy002’, and bacterial growth was determined at 0, 6, and 12 days post-inoculation. Error bars represent standard error of the mean from four pseudo-biological replicates. Asterisks indicate results of Student’s *t*-test between selected sample and wild-type; **(*P* < 0.01), ***(*P* < 0.001). The experiment was conducted three times with similar results. **(B)** *P*sa BV1, BV1 LPS knock-out mutant (Δ*LPS*), two independent BV1 knock-outs complemented LPS from BV3 (+LPS_BV3), and two independent BV1 revertant-to-knock-out LPS (-LPSrev) were flood-inoculated at ∼10^6^ CFU/mL on *A. chinensis* var. *chinensis* ‘Hort16A’, and bacterial growth was determined at 0, 6, and 12 days post-inoculation. Error bars represent standard error of the mean from four replicates. Asterisks indicate results of Student’s *t*-test between selected sample and wildtype (*Psa* V13); ***(*P* < 0.001). The experiment was conducted twice with similar results.

### The TET operon from *P. syringae* isolates is highly variable but correlates with LPS

Having identified the TET operon in *P. syringae* for the first time we set out to analyse this operon in the *P. syringae* species complex and closely related plant associated *Pseudomonas* species. In *P. syringae*, the composition of the CPA backbone (D- and/or L-Rha), the number of sugars in the repeat, and the bonds linking the sugars within the oligosaccharide, vary between different *P. syringae* isolates (Ovod *et al.*, 1997a; Zdorovenko and Zdorovenko, 2010). BLASTx and Conserved Domain Database (CDD) searches identified the CPA locus in other *P. syringae* pathovars and epiphytic isolates. We also included several *P. viridiflava* isolates (See Supplementary Table 1 for details of all isolates analysed).

Comparison of these loci revealed variation in the sugar biosynthetic pathways comprising the CPA locus, including the TET operon. In some *P. syringae* isolates, the GT and O-methyltransferase genes were fused. The TET operon is the only one consistently present at the CPA locus, and it showed a high degree of sequence variation between *P. syringae* pathovars and isolates. Variation in CPA sugar biosynthetic pathways and the sequence divergence of the TET operon made the identification of this region in *P. syringae* challenging, but as noted previously, the CPA locus was consistently located adjacent to genes encoding a chemotactic receptor, DnaJ molecular chaperone and a tRNA-Met (Figure 1A). A gene encoding an integrase was often located adjacent to the tRNA-Met in an orientation reminiscent of an integron (Domingues *et al.*, 2012; Gillings, 2014). Given that tRNAs are often sites for integration of mobile elements, this might explain the modular and variable nature of the CPA locus in *P. syringae*. However, a canonical integron structure was not found using programs such as integron finder (Cury *et al.*, 2016).

Phylogenetic analysis of the concatenated proteins from the TET operon generated a tree with five main clades (including the two clades identified in *Psa* and *Pfm*), each representing different sets of alleles of this operon from the *P. syringae* and *P. viridiflava* species complexes (Figure 5A). As observed with *Psa* and *Pfm* isolates, the topology of the tree made from these sequences is very different from the corresponding tree generated from five core genes from the respective genomes (Supplementary Figure S1). This indicates that the TET operon has an evolutionary history driven by recombination or horizontal gene transfer that is distinct from the evolution of the core genome.

**Figure 5.**
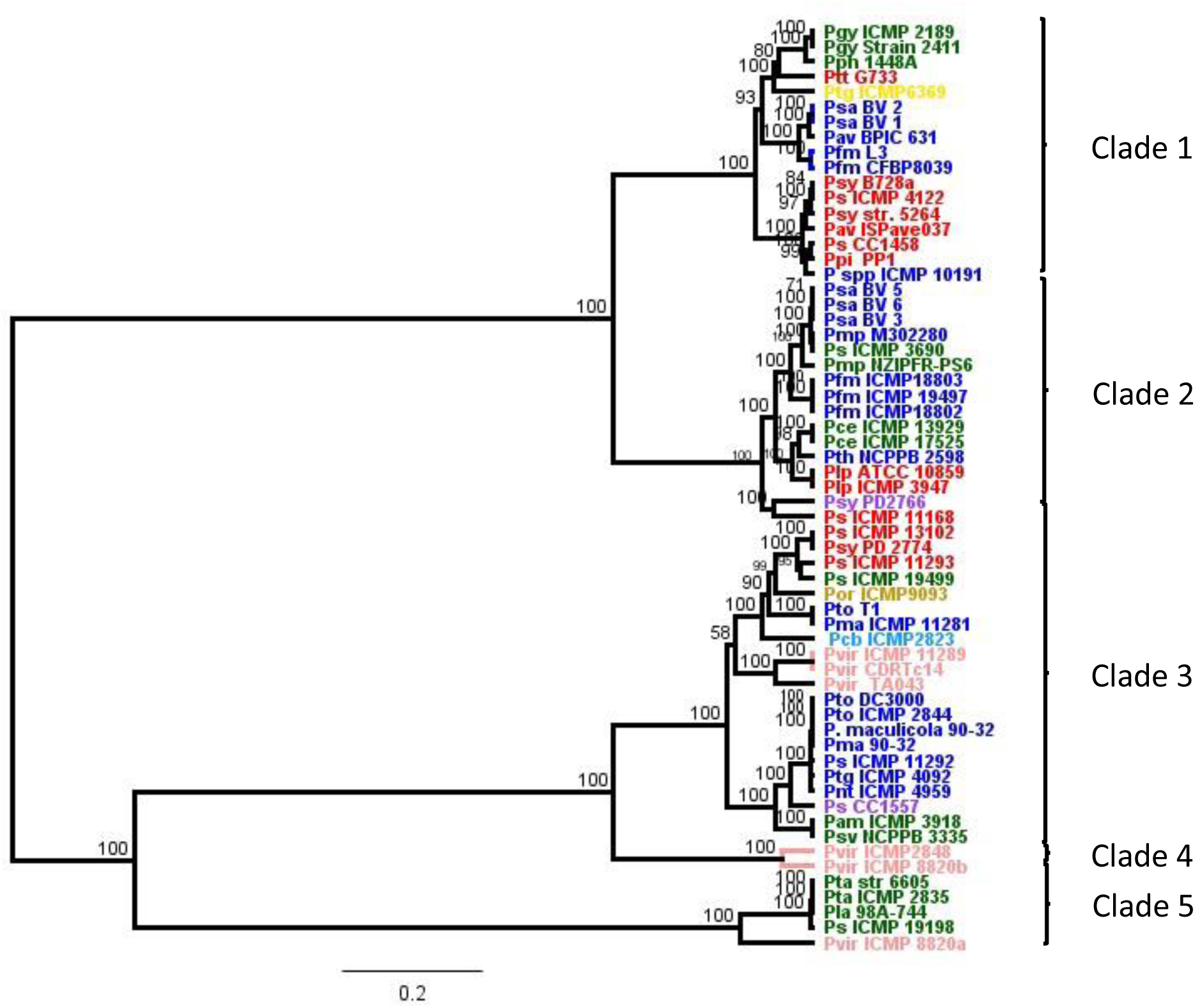
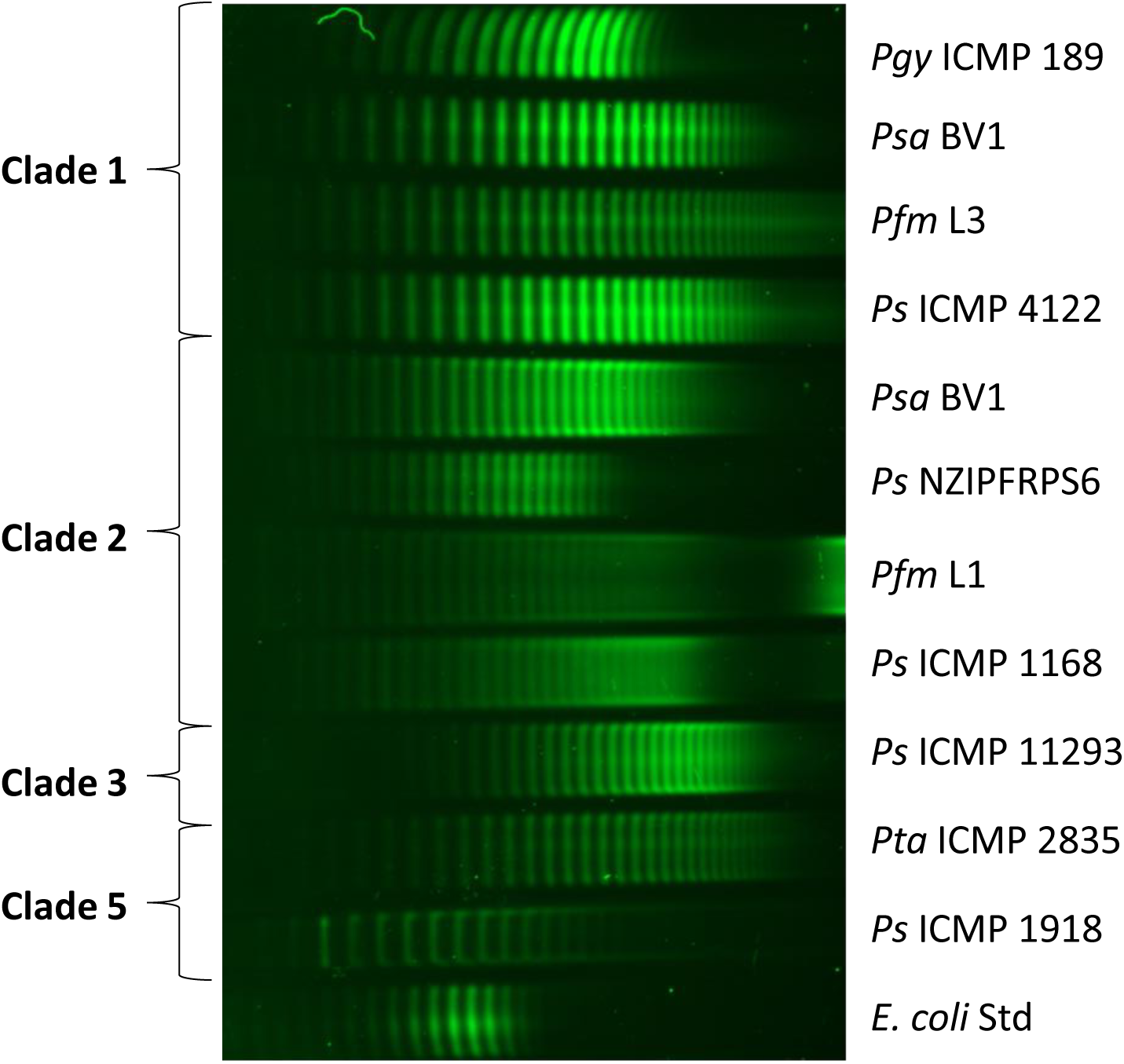
**(A)** Phylogenetic comparison of TET operons from various *Pseudomonas syringae* strains. Amino acid sequences from three proteins of the CPA locus (Wzt, WbdD and WbdA) were aligned in Geneious v10 and neighbour-joining phylogenetic trees built for their TET operons. Strains were grouped into numbered clades as indicated. Strains are coloured according to their core genome phylogenies as previously described (Dillon *et al.*, 2019b): phylogroup 1 blue; phylogroup 2 red; phylogroup 3 green; phylogroup 4 orange; phylogroup 5 purple; phylogroup 6 yellow; phylogroup 7 pink; phylogroup 9 grey; phylogroup 10 mauve; phylogroup 11 chocolate; phylogroup 13 claret. **(B)** LPS profiles of *Pseudomonas syringae* representing phylogenetic clades for the TET operon. Pro-Q™ Emerald 300 staining of lipopolysaccharide (LPS) preparations on a 4–12% gradient Tris-glycine SDS gel from representatives of *P. syringae* strains marked with asterisks from Supplementary Table 1 (top to bottom, lanes 1–11), with *Escherichia coli* control (Std, lane 12). Banding patterns were grouped into clades on their designations from Figure 7. A total of 2–15 μL LPS extract for each sample (normalized by trial runs) was applied per lane.

Since genetic variation in the CPA between even closely related isolates of *Psa* and *Pfm* is reflected in different LPS profiles, we performed a broader analysis of LPS profiles across different clades of *P. syringae* (Figure 5A). Eleven isolates of *P. syringae* were chosen, representing four TET operon phylogenetic clades. The banding patterns were different for all isolates, but similarity within clades was observed. Notably, the spacing between bands was clearly similar within clades, but varied between them (Figure 5B). Interestingly, the range of molecular weights of different LPS molecules within clades was highly variable, suggesting the efficiency of the chain terminating enzyme (WbdD) varies within a clade. While considerable variability has been demonstrated in *P. syringae* CPA, the functional relevance of this is less well understood.

### Correlation between tailocin sensitivity and TET operon variability

Like phages, many R-type syringacins (tailocins) use LPS as a receptor. Recently the R-syringacin killing and sensitivity spectra for a diverse range of *P. syringae* isolates were characterised (Baltrus *et al.*, 2019). A feature of both spectra is that dendrograms generated from the killing and sensitivity matrices do not match MLST phylogenetic trees (Baltrus *et al.*, 2019). Since LPS is a likely receptor for R-syringacins, we hypothesised that there might be a correlation between the patterns of variation in tailocin sensitivity within *P. syringae* isolates and the genetic variation in the TET operon responsible for the structure of the CPA backbone.

To test this hypothesis, we generated a UPGMA phylogenetic tree from the TET operons from 29 of the isolates tested for tailocin killing and sensitivity by Baltrus *et al.* (2019). This was compared to a hierarchical clustering tree generated from the sensitivity profile of the same isolates using a tanglegram (Figure 6A). Tanglegrams compare phylogenetic trees to determine how similar they are and correlation coefficients can be calculated from the similarity between different pairs of trees (Galili, 2015). The correlation coefficient between the sensitivity and TET operon trees was 0.987, indicating a high degree of congruence (Supplementary Figure S6). Both trees comprise two main clades, and there was a 100% correlation among the isolates between the clades in both trees. In contrast, tanglegrams between either the sensitivity profile or the TET operon and an MLST phylogenetic tree had low correlation coefficients of 0.011 and -0.003, respectively, indicating poor congruence (Figure 6B-C; Supplementary Figure S5). These results suggest a strong relationship between tailocin sensitivity and the structure of the CPA oligosaccharide of LPS in *P. syringae*. They provide further evidence that LPS is a tailocin receptor in *P. syringae*, explaining the complex host range observed for these proteins (Baltrus *et al.*, 2019) and providing a plausible explanation for the diversification pattern of these loci.

**Figure 6.**
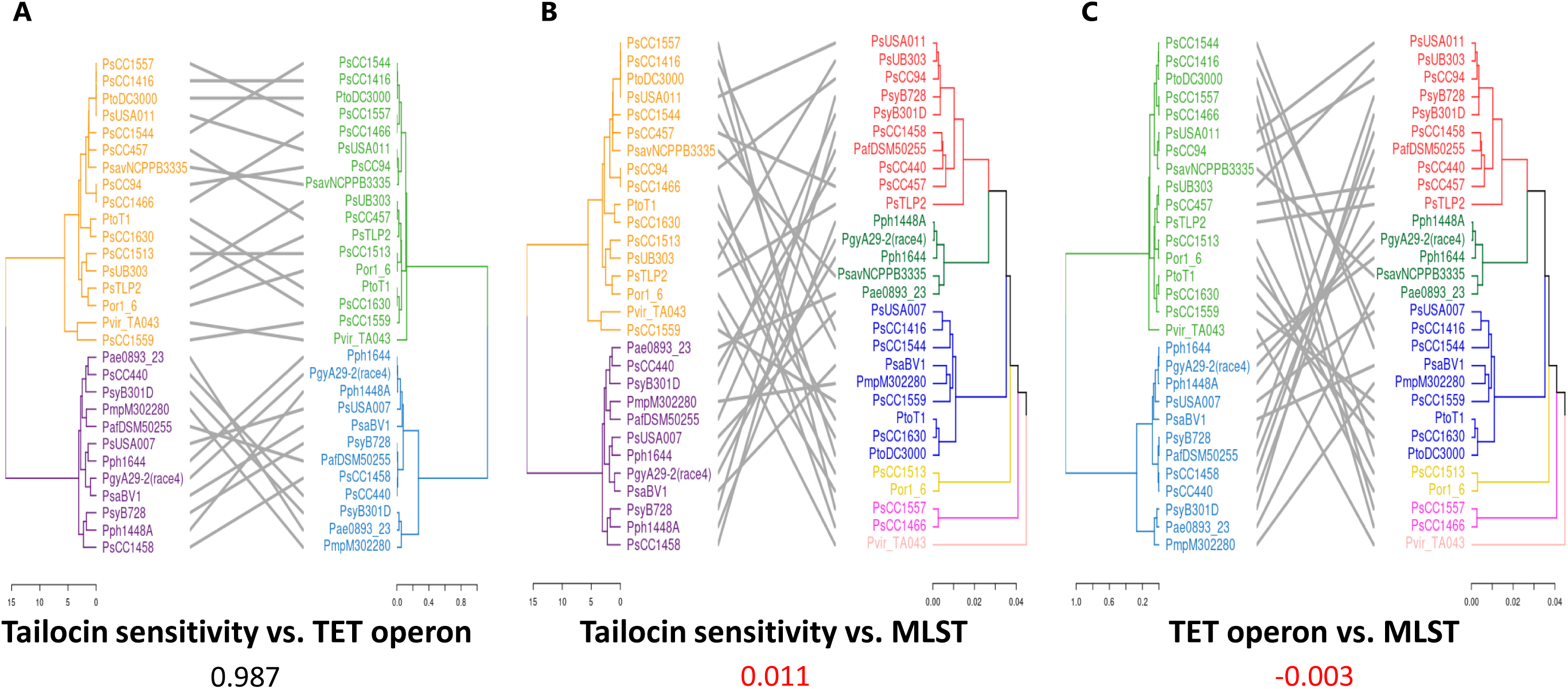
Tanglegram comparison of tailocin sensitivity, TET operon, and core gene MLST phylogenies. **(A)** Tanglegram of sensitivity matrix phylogeny (left) and TET operon phylogeny (right). **(B)** Tanglegram of sensitivity matrix phylogeny (left) and core gene MLST phylogeny (right). **(C)** Tanglegram of TET operon phylogeny (left) and core gene MLST phylogeny (right). Hierarchical clustering analysis using the ward.D2 method was used to build a tailocin sensitivity phylogeny from the sensitivity matrix provided in Baltrus et al. (2019), which is transposed from a syringacin killing matrix. The TET operon phylogeny was produced by using the TET operon genome sequences of selected strains, assembled into a phylogeny using the UPGMA clustering method, as was the MLST phylogeny. The co-phenetic entanglement coefficients for each comparison are indicated below each panel title. Axes in each panel represent Euclidean distance. Colours represent broad clade groupings, with the cut-offs being Euclidean distances of 10, 0.5, and 0.02 for the sensitivity, TET, and MLST phylogenies, respectively. This Euclidean cut-off for the MLST branches allows representative separation by phylogroup for the strains used.

## DISCUSSION

LPS is a highly complex macromolecule located on the surface of the bacterial cell wall that interfaces with the environment and other organisms. It is often used as a receptor or ligand for recognition by other bacteria, phages, bacteriocins and the defence system of host organisms. The modular structure can be used to generate structural variability, particularly in the OPS, which can help bacteria avoid parasitic organisms, toxins and host defences (King *et al.*, 2009). The structure and function of LPS has been well explored in human and animal bacterial pathogens where it has been shown that LPS interacts directly with the host immune system and hence immunity is a strong driver for genetic variation at loci coding for LPS (Lerouge and Vanderleyden, 2002; Maldonado *et al.*, 2016).

In contrast, the role of LPS in plant-pathogenic and epiphytic bacteria is less clear. In this study, we have identified the CPA locus in *Psa* and shown it is responsible for the biosynthesis of the predominant OPS in *P. syringae*. The locus is highly modular with operons predicted to be involved in the biosynthesis of various sugars, although only those for L-Rha and D-Rha have been convincingly identified through similarity. The TET operon that codes for proteins involved in the polymerisation and transport of the backbone oligosaccharide through the inner membrane was also identified for the first time in *P. syringae.* This was made possible by the use of LPS screens of our transposon library (Mesarich *et al.* 2017). This operon was present in the majority of *P. syringae* and *P. viridiflava* CPA loci examined. Phylogenetic analysis of three of the proteins in the TET operon revealed five distinct clades, which most likely reflect the different unit structures of the CPA Rha-backbone oligosaccharide. There are two elements that contribute to the structural variability of the CPA in *P. syringae*. One is the nature of the backbone oligosaccharide, which can be tri- or tetra-saccharides of either D-Rha, L-Rha or both, linked by α1-2, α1-3, or β1-4 bonds (Ovod *et al.*, 1997a). The other is the side-chain decoration, which is usually a single sugar residue attached by a variety of different linkages. In addition, acetylation is a common back-bone modification. Unfortunately it is not possible to directly correlate the structural information accurately with the clades resolved in Figure 5 because the isolates for which structural LPS information is known are not those that have been sequenced (Zdorovenko and Zdorovenko, 2010).

The mechanism by which the different operons within the CPA locus are shuffled between *P. syringae* isolates to generate the observed structural variability could occur via homologous recombination or insertion via an integron-like cassette. The structure of the CPA locus in *Psa* BV3 resembles that of an integron, although an intact phage-like integrase adjacent to the tRNA-Met was not present in all other *P. syringae* isolates. In addition, a classic integron promoter and *attC* insertion sequences were not found using programs such as integron finder. The presence of operons from *P. viridiflava* in the phylogenetic trees suggest that the mechanism of recombination between isolates is not exclusive to the *P. syringae* species complex, but also includes other closely related species such as *P. viridiflava* that share the same ecological (plant-associated) niche.

The variation at the TET locus is also reminiscent of that at the capsule locus within the *Klebsiella pneumoniae* clonal group 258 (Wyres *et al.*, 2015). In this case it has been shown that extensive intra-clonal variation at the capsule polysaccharide synthesis locus is mediated by large-scale recombination events. This has been responsible for generation of the 78 immunologically distinct capsule variants which have been described in *K. pneumoniae* (Wyres *et al.*, 2015).

Tailocins are a subset of bacteriocins derived from phage tail proteins that have been co-opted by host bacteria as a means for killing closely related bacterial isolates occupying the same ecological niche (Ghequire and De Mot, 2015). What has been demonstrated recently within *P. syringae* is the depth and complexity of both the killing and sensitivity spectra generated from a wide range of isolates (Baltrus *et al.*, 2019). While it is clear that LPS is a receptor for some tailocins, a link between the sensitivity to tailocins and the structural variation in LPS has not been demonstrated until now. Our results indicate a strong relationship, suggesting that killing by tailocins (and by inference infection by phage) may drive diversity in LPS structure. It is interesting to note that the genetic variation driving these differences appears in both cases to be mediated by localised recombination events (Baltrus *et al.*, 2019).

As has been reported previously for other *P. syringae* LPS mutants (Kutschera *et al.*, 2019), those in both the *Psa* BV3 and BV1 backgrounds displayed significantly reduced *in planta* growth compared to their wild-type strains at both 6- and 12-days post-inoculation (Figure 4). The inability of the LPS knockout isolates to remain suspended in solution may also affect the ability of these mutants to move *in planta* and could be responsible for reduced pathogenicity. Although mutants lacking LPS have been shown to be highly resistant to particular bacteriophages and/or tailocins, the loss of pathogenicity probably ensures that strain completely lacking LPS are strongly selected against *in planta* (Kandel *et al.*, 2019). This means that gene replacements at this locus should be favoured over simple gene deletion events.

There did not appear to be any obvious geographical bias for any of the CPA clades. While this is unlikely to be an exhaustive identification of TET operon variants, a significant proportion of available *P. syringae* whole genome sequences were analysed in this study. Since members of four clades were isolated recently from kiwifruit in New Zealand, the TET operon alleles identified are likely ubiquitous among *P. syringae* populations. These findings, firstly that LPS is a trait conferred by a predictable locus, and secondly that LPS is required for pathogenicity, offer intriguing possibilities for a durable disease resistance program using bacteriophage/tailocin-mediated control. While the locus is open to recombination, the discovery of the population of TET operon alleles present on a plant species may facilitate a comprehensive mixture of bacteriophage/tailocin that covers the most common alleles to be deployed to increase efficacy and durability of this treatment by circumventing escape through recombination at the TET operon locus.

## EXPERIMENTAL PROCEDURES

### Bacterial isolates used in this study

Metadata for the bacterial isolates and genomes used in this study are listed in Supplementary Table 1. Lysogeny broth (LB), and M9 minimal media (M9) (Sambrook and Russell, 2001) were supplemented with agar 1.5% (w/v) and 50 µg/mL kanamycin (Km) where required. *Escherichia coli* and *Pseudomonas* isolates were routinely cultured at 37°C and 28°C, respectively. All kits and reagents were used, except where specified, in accordance with the manufacturer’s instructions.

### Bioinformatics

Bioinformatics analysis was carried out using the Geneious R10 platform (www.geneious.com). For unannotated genomes, the RAST pipeline was used for gene calling (Aziz *et al.*, 2008). GenBank searches were carried out using BLAST (Boratyn *et al.*, 2013). Sequences were searched for *attI* and *attC* motifs using Integron Finder https://github.com/gem-pasteur/Integron_Finder.

### LPS purification

For small-scale preparations of LPS, the hot-phenol method was used (Westphal and Jann, 1965). Overnight bacterial cultures were resuspended in PBS (500 μL) to an A_600_ of between 2 and 5. An equal volume of Tris-saturated phenol (Sigma-Aldrich, Mo, USA, P4557) was added and the sample was incubated at 68°C for 90 min with intermittent vortexing. The supernatant was sequentially treated with DNase I (37°C) and RNase (65°C), each for 1 h. The samples were treated with phenol/chloroform/isoamyl alcohol, followed by chloroform/isoamyl alcohol and stored at -20°C. To generate antigen used for immunization and polyclonal antibody generation, large-scale extraction of LPS from *Psa* BV1 and BV3, and *Pfm* L1 was performed using the hot-phenol method of Maier and Brill (1978). Cells were harvested in late exponential phase, collected by centrifugation and freeze-dried for storage at -20°C.

LPS were resolved using 4–12% premade NuPAGE™ Bis-Tris SDS gels on a Novex system with MOPS as the running buffer. Gels were stained using the Pro- Q™ Emerald 300 Lipopolysaccharide Gel Stain Kit (Molecular Probes, OR, USA). For Western blotting, gels were transferred to a polyvinylidene difluoride (PDVF) membrane (Millipore, Burlington, MA). The anti-Psa antibodies (see below) were used at 1:5,000 dilution and the Anti-Rabbit IgG-Peroxidase antibody (Sigma-Aldrich, St Louis, MO) was used at 1:10,000 dilution. Chemiluminescence was developed using the Clarity Max Western ECL (Bio-Rad, Hercules, CA).

### Production of antibodies to *Psa* biovars

Heat-killed cells from *Psa* BV1 (ICMP 9617), *Psa* BV3 (ICMP 18884), and *Pfm* L1 (ICMP 18803) in Freund’s complete adjuvant were each used to individually immunise two rabbits by subcutaneous injection of 10^8^ cells. Rabbits were rested for 4 weeks and immunised every 2 weeks by three further injections. Blood was collected 1 week after the last injection, and rabbit IgG was purified from serum using ammonium sulphate fractionation and protein-A sepharose chromatography (Ey *et al.*, 1978). These polyclonal antibodies were further purified by affinity chromatography through matrices attached to LPS extracted from the above isolates not used for immunisation in each of the three cases. The aim of this step was to remove antibodies that bound to antigens in common with other isolates, to give an isolate-specific polyclonal antibody preparation. For example to purify antibodies specific for *Psa* BV3, IgG were passed through affinity columns with LPS from *Psa* BV1 and *Pfm* L1. For preparation of the matrices LPS was dissolved in buffer and oxidised using periodate. Excess periodate was removed by gel filtration using a PD10 column. Oxidised LPS was reacted with biotin hydrazide to biotinylate the LPS as per method sheet 28020 (Thermo Fischer Scientific). LPS matrices for affinity chromatography were prepared by attachment of biotin-LPS to streptavidin-labelled sepharose columns. Biovar-specific LPS antibodies were purified by standard affinity chromatography as described by the manufacturer (GE Life Sciences, PA USA).

### LPS locus knock-out and knock-in

To generate the LPS TET operon (*wzt* – *wbdA*; ∼5.3 kb) deletion in *Psa* BV1 (ICMP 9617), a modified pK18mobsacB vector-based method was utilized (Kvitko and Collmer, 2011). DNA fragments containing the upstream (∼1 kb) and downstream (∼1 kb) regions of the LPS knock-out region were amplified using PCR with primer pairs PsaJ_LPS-KO_UP-F/PsaJ_LPS-KO_UP-R and PsaJ_LPS-KO_DN- F/PsaJ_LPS-KO_DN-R, respectively, with the UP-R and DN-F primers carrying added *Xba*I restriction enzyme sites (Supplementary Table 2). Each PCR fragment was gel-purified using an EZNA gel extraction kit (Custom Science, Auckland, NZ) and digested with *Xba*I, re-purified with an EZNA PCR product purification kit and ligated (to the *Xba*I-cut overhangs) to form a 2 kb KO fragment. The 2 kb fragment containing both the upstream and downstream fragments for the KO region was then re-amplified by PCR using primers PsaJ_LPS-KO_UP-F and PsaJ_LPS-KO_DN-R. This PCR product was cloned into the *Eco53*kI blunt-end restriction enzyme site of pK18mobsacB that had first been mutagenized to remove the non-MCS Eco53kI site to generate pK18BΔE (Schafer *et al.*, 1994). The pK18BΔE vector carrying the ∼2 kb KO fragment, called pΔ*LPS*, was transformed into *E. coli* DH5α, plated on X- gal/IPTG/kanamycin LB agar for blue/white selection. Positive transformants were confirmed by Sanger sequencing (Macrogen, Seoul, South Korea). *Psa* BV1 was electroporated with the pΔ*LPS* construct and transformants were selected on LB agar with nitrofurantoin, cephalexin and kanamycin. Selected colonies were subsequently streaked onto LB agar containing 10% (w/v) sucrose to counter-select plasmid integration, forcing removal of the *sacB* gene, resulting in either revertants to wild-type or knock-outs. KOs were confirmed using colony PCR with primers PsaJ_LPS-KO_Check-F and PsaJ_LPS-KO_Check-R, with PCR products sent for Sanger sequencing with the nested cloning PsaJ_ LPS -KO_UP-F and PsaJ_ LPS - KO_DN-R primers, as well as using internal gene-specific primers. Selected colonies were also plated on kanamycin-containing medium to confirm loss of the plasmid backbone containing kanamycin resistance and the *sacB* gene.

To clone the knock-in (KI) construct of the *Psa* BV3 LPS TET operon for complementation of the *Psa* BV1 LPS TET operon knockout (*Psa* BV1 Δ*LPS*), the golden gate cloning system was used (Engler *et al.*, 2009). Firstly, the destination vector was generated by restriction digestion using *Ase*I and *Nhe*I of pK18mobsacB to remove the multiple cloning site (MCS). This was replaced with a golden gate-compatible (*Bsa*I restriction enzyme site-flanked) MCS from pICH86988 using In- Fusion cloning according to kit instructions (Takara Bio, USA) to generate a golden gate cloning-compatible version of the pK18mobsacB vector; pK18B-GG. The pK18B-GG vector was transformed into *E. coli* DH5α, plated on kanamycin LB agar and screened by *Bsa*I restriction digest. Positive transformants were confirmed by Sanger sequencing (Macrogen, South Korea). Entry-level vectors for the LPS-KI construct were generated for the LPS region from *Psa* BV3 (ICMP 18884), divided into four modules (modules 1–4), with overlap-fusions of the upstream (5’) and downstream (3’) regions of the *Psa* BV1 Δ*LPS* (for modules 1 and 4, respectively, with modules made by overlap PCR) and an additional module 5 for the downstream region of the *Psa* BV1 Δ*LPS* by PCR-amplification. These were cloned into pICH41021 with the cloning primers (Supplementary Table 2) designed to synonymously mutate internal *Bsa*I, *Eco53k*I, and *Pst*I restriction enzyme sites (Supplementary Figure S2). These five LPS-KI modules in pICH41021 were then single-pot cloned (by golden gate assembly) into destination pK18B-GG to generate the LPS-KI vector: pLPSbv3. The pLPSbv3 vector was transformed into *E. coli* DH5α, plated on kanamycin LB agar and screened by nested colony PCR for correct assembly. Positive transformants were confirmed by Sanger sequencing (Macrogen, South Korea). *Psa* BV1 Δ*LPS* was transformed with pLPSbv3 and transformants were selected on LB agar with nitrofurantoin, cephalexin and kanamycin. Selected colonies were subsequently streaked onto LB agar containing 10% (w/v) sucrose to counter-select plasmid integration, resulting in revertants to knock-out or LPS- complemented mutants. Complementation knock-in mutants were screened using colony PCR with primers for PsaJ_LPS-KO_Check-F and PsaJ_LPS-KO_Check-R, as well as gene-specific primers and sent for Sanger sequencing. Complemented mutants (Δ*LPS +LPS_BV3*) and non-complemented knock-out revertants (Δ*LPS- LPSrev*) were further confirmed by plating on kanamycin-containing agar to confirm loss of the *sacB* gene.

### *In planta* growth of *Psa*

*Psa* infection assays were based on those published previously with some modifications (McAtee *et al.*, 2018). *A. chinensis* Planch. var. *chinensis* ‘Zesy002’ or ‘Hort16A’ plantlets, grown from axillary buds on Murashige and Skoog rooting medium without antibiotics in sterile 400-mL plastic tubs (pottles) with three plantlets per pottle, were purchased (Multiflora, Auckland, NZ). Plantlets were kept at 20°C under Gro-Lux fluorescent lights under long-day conditions (16 h light: 8 h dark) and used when the plantlets were between 10–14 weeks old. Overnight liquid cultures of wild-type or mutant strains of *Psa* were pelleted at 6000 *g*, re-suspended in 500 mL of 10 mM MgSO_4_ to an A_600_ = 0.05 (∼107 CFU/mL, determined by plating). Surfactant Silwet™ L-77 (Lehle Seeds, Round Rock, USA) was added to the inoculum at 0.0025% (v/v) to facilitate leaf wetting. Pottles of ‘Zesy002’ or ‘Hort16A’ plantlets were filled with the inoculum, submerging the plantlets for 3 min, drained, sealed, and then incubated under previously described plant growth conditions. To assess *in planta* growth, leaf samples of four leaf discs per replicate, removed with a 1-cm diameter cork-borer, were taken at 2 h (day 0), day 6, and day 12 post-inoculation. All four pseudo-biological replicates per treatment were taken from the same pottle. To estimate *Psa* growth inside the plant, the leaf discs were surface sterilized, placed in Eppendorf tubes containing three sterile stainless-steel ball bearings and 350 μL 10 mM MgSO_4_, and macerated in a Storm 24 Bullet Blender (Next Advance, NY, USA) for two bursts of 1 min each at maximum speed. A 10-fold dilution series of the leaf homogenates was made in sterile 10 mM MgSO_4_ until a dilution of 10-8 and each dilution was plated as 10 μL droplets on LB agar supplemented with nitrofurantoin and cephalexin. After 2 days of incubation at 20°C, the CFU per cm^2^ of leaf area was ascertained from dilutions.

### Hierarchical clustering and dendrogram analysis

Hierarchical cluster analysis was conducted in R (R-CoreTeam, 2018) using the ward.D2 method, as per the clustering analysis in Baltrus et al. (2019). The ‘phylogram’ vignette was used to convert hierarchical clusters into dendrograms (Paradis and Schliep, 2018). The tanglegram function in the ‘dendextend’ package was used to visually compare dendrograms by connecting matching leaf node labels with lines (Galili, 2015). The ladder function was used to untangle dendrograms by rotating the tree branches at their nodes without altering their topology, allowing for clear visualization (Galili, 2015). The cor.dendlist function was used to calculate the cophenetic correlation coefficient for the compared dendrograms (Galili, 2015). Cophenetic correlation values (correlation coefficients) range from -1 to 1, with values close to 0 indicating that the compared dendrograms are not statistically similar. The resulting cophenetic correlation matrix was then visualized using the ‘corrplot’ package (Wei and Simko, 2017).

## ACKNOWLEDGEMENTS

This work was funded (including a post-doctoral fellowship to JJ) by the Bio- protection Research Centre (Tertiary Education Commission). LMH would like to thank Zespri International for an MSc scholarship.

We would like to thank Rick Broadhurst (AgResearch, Ruakura, New Zealand) for rabbit immunology. We would also like to thank Jo Bowen (PFR) and Iain Hay (UoA) for critically reading the manuscript.

## FIGURE LEGENDS

**Supplementary Figure S1.**
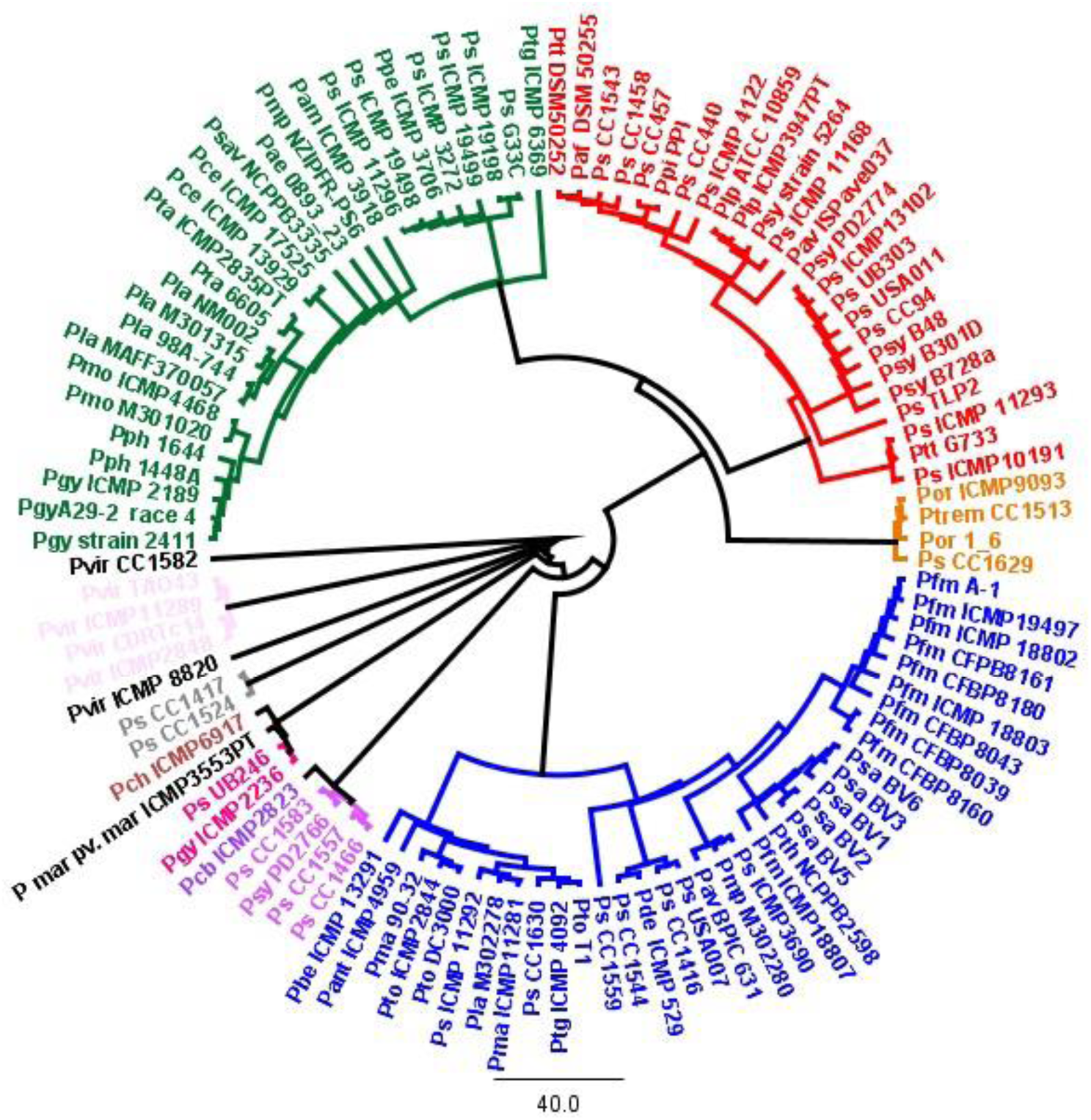
MLST genome phylogenetic tree for *Pseudomonas syringae* and *Pseudomonas viridiflava* strains. Five genes (*gyrB, ropD, gapA, pgi* and *acnB*) from selected representatives of *P. syringae* phylogroups were concatenated and aligned in Geneious v11 and neighbour-joining phylogenetic trees built. Strains grouped as expected into their phylogroups and are coloured according to their core genome phylogenies as previously described (Dillon *et al.*, 2019b): phylogroup 1 blue; phylogroup 2 red; phylogroup 3 green; phylogroup 4 orange; phylogroup 5 purple; phylogroup 6 yellow; phylogroup 7 pink; phylogroup 9 grey; phylogroup 10 mauve; phylogroup 11 chocolate; phylogroup 13 claret.

**Supplementary Figure S2.**
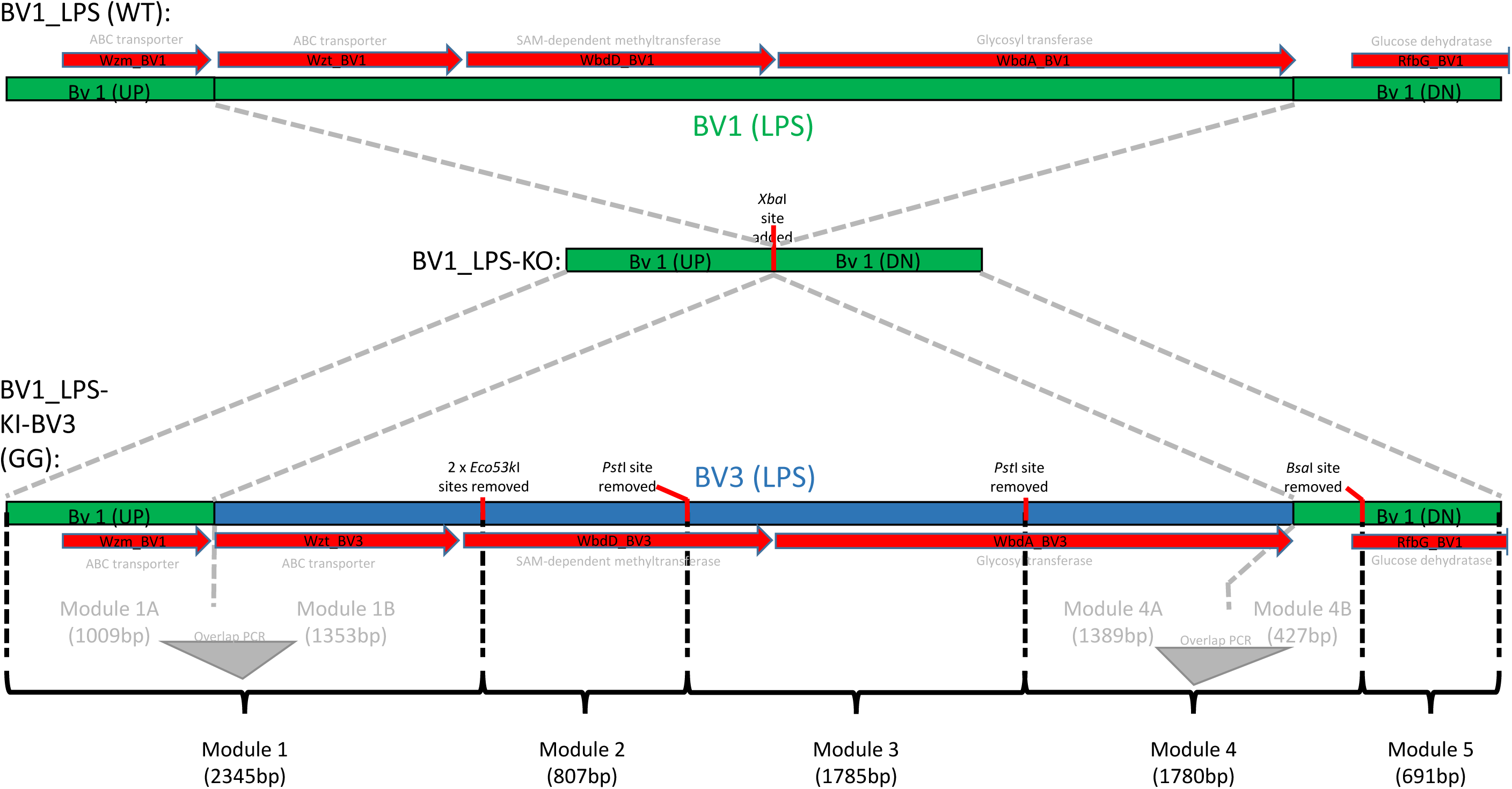
Schematic for knock-out and knock-in swap of the TET locus in *Pseudomonas syringae* pv. *actinidiae* (*Psa*) BV1. One-kilobase flanking regions upstream (UP) and downstream (DN) of the polymorphic region of the TET operon in wild-type (WT) *Psa* BV1 (green; encoding Wzt, WbdD, and WbdA proteins in red) were PCR-cloned and fused at a primer-introduced internal *Xba*I site and used in pK18mobsacB vector-mediated knock-out of the BV1 TET locus to generate BV1_LPS-KO. To generate the BV1-to-BV3 LPS swap (knock-in complementation), a pK18mobsacB vector-mediated method was also used. First, *Eco53k*I, *Bsa*I and *Pst*I sites were removed during PCR (primer-driven) to facilitate blunt-end cloning of modules, golden gate (GG) cloning, and evasion of the BV1-encoded restriction enzyme, respectively. The knock-in pK18mobsacB vector was built (GG assembly) from five modules with sizes indicated. Modules 2, 3, and 5 were PCR-amplified and blunt-end cloned into pICH41021 shuttle vector directly. Modules 1 and 4 were first cloned as two parts (A and B) from their respective *Psa* BVs and then fused by overlap PCR mediated by primer sequences designed to introduce overlap regions during amplification of part A and B. Following overlap PCR, modules 1 and 4 were blunt-end cloned into the pICH41021 shuttle vector. The modules were then assembled into the final pK18mobsacB knock-in construct. The BV1_LPS-KO was then subsequently complemented with TET operon from BV3 (blue).

**Supplementary Figure S3.**
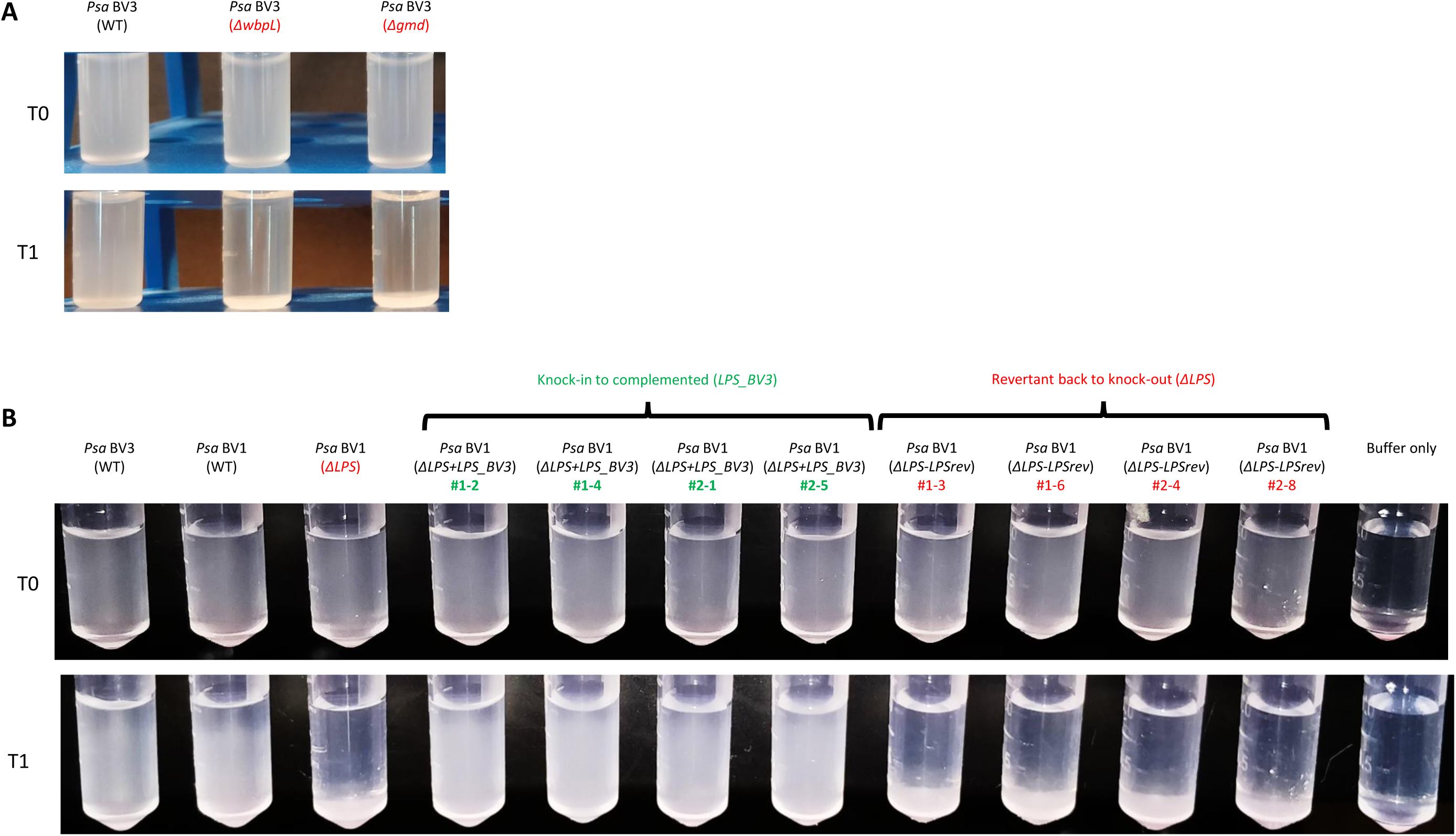
Bacterial cell sedimentation for *Pseudomonas syringae* pv. *actinidiae* (*Psa*) in inoculation buffer. (A) *Psa* BV3 and two R-LPS mutants, Δ*wbpL* and Δ*gmd* (Mesarich *et al.*, 2017) were grown overnight in LB liquid culture, pelleted by centrifugation, resuspended in 10 mM MgSO_4_ buffer, and normalized to OD_600_ of 0.4. (B) *Psa* BV3 and BV1 (wild-type; WT), *Psa* BV1 LPS knock-out (Δ*LPS*), four BV1-to-BV3 TET operon-complemented isolates (Δ*LPS+LPS_BV3*: #1-2, #1-4, #2-1, and #2-5; green), and four BV1 revertant to knock-out isolates (Δ*LPS-LPSrev*: #1-3, #1-6, #2-4, and #2-8; red) were grown overnight in LB liquid culture, pelleted by centrifugation, resuspended in 10 mM MgSO_4_ buffer, and normalized to OD_600_ of 0.5. Photographs of cell suspensions in both (A) and (B) were taken immediately after resuspension in buffer (T0) and after an hour standing at room temperature (T1).

**Supplementary Figure S4.**
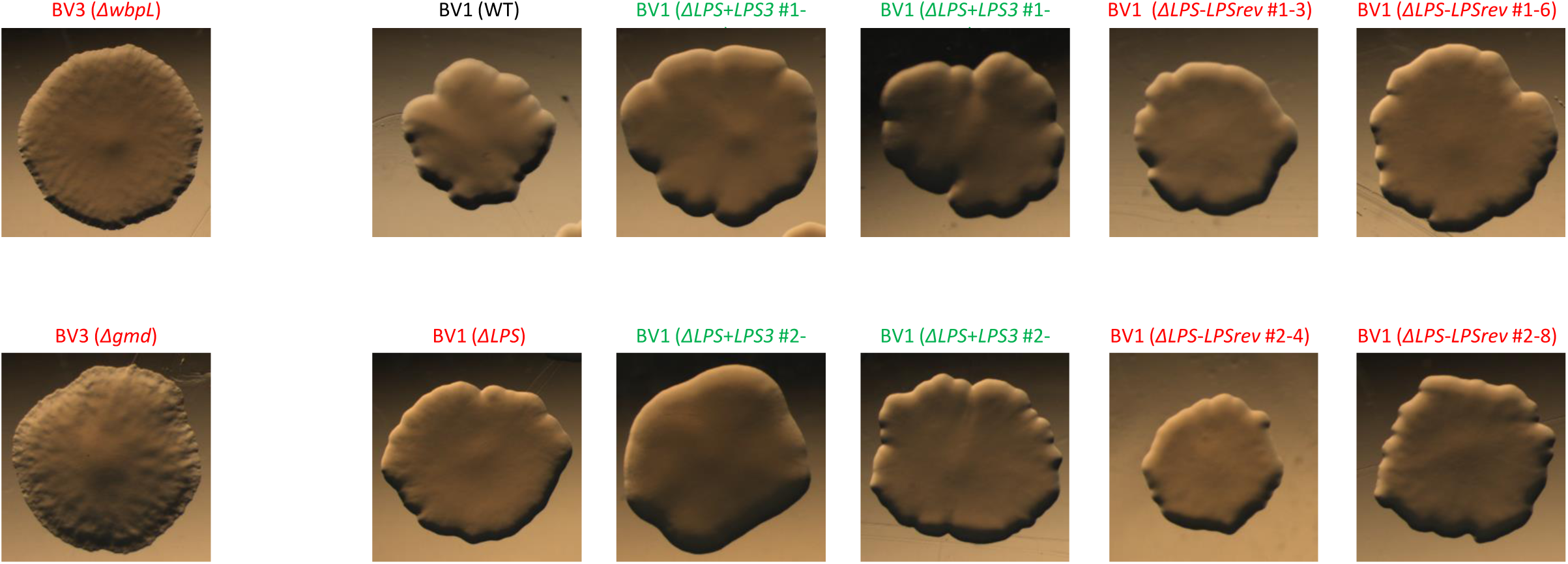
Colony morphologies for *Pseudomonas syringae* pv. *actinidiae* (*Psa*) BV1 mutant and complemented strains. *Psa* BV3 mutants (Δ*wbpL* or Δ*gmd*) (Mesarich *et al.*, 2017) and BV1 (wild-type; WT), *Psa* BV1 LPS knock-out (Δ*LPS*), four BV1-to-BV3 TET operon complemented isolates (Δ*LPS+LPS_BV3*: #1- 2, #1-4, #2-1, and #2-5; green), and four BV1 revertant to knock-out isolates (Δ*LPS- LPSrev*: #1-3, #1-6, #2-4, and #2-8; red) were grown on LB agar plates, for 3 days and photographs of colonies were taken under a stereomicroscope.

**Supplementary Figure S5.**
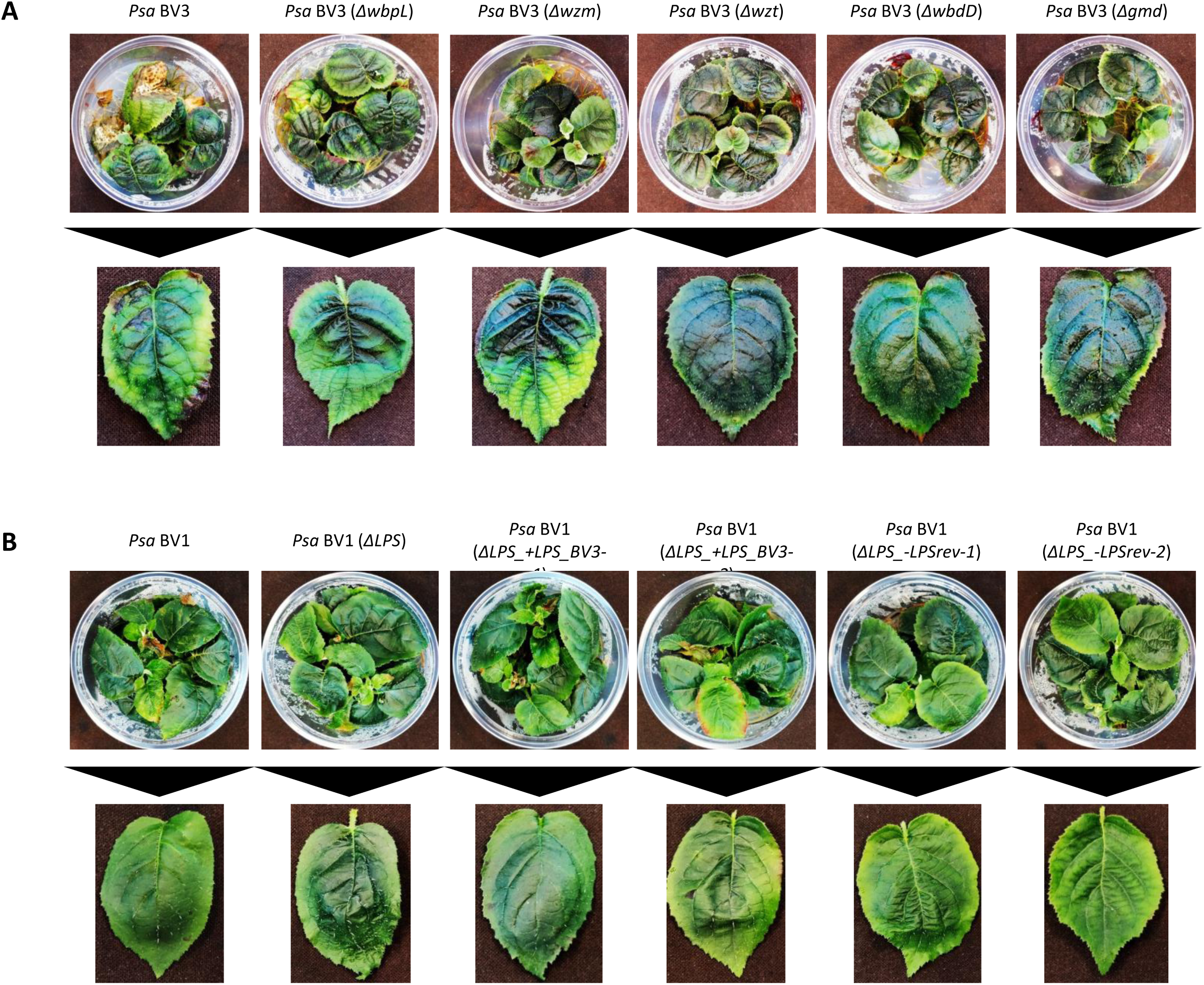
The CPA TET operon is required for symptom development of *Pseudomonas syringae* pv. *actinidiae* (*Psa*) in host plants. **(A)** *P*sa BV3 or BV3 R-LPS mutants ((Mesarich *et al.*, 2017)) were flood-inoculated at ∼10^6^ CFU/mL on *Actinidia chinensis* var. *chinensis* ‘Zesy002’, and photographs of symptom development in pottles and representative leaves taken at 50 days post-inoculation. **(B)** *P*sa BV1, BV1 LPS knock-out mutant (Δ*LPS*), or two independent BV1 knock-outs complemented LPS from BV3 (+*LPS_BV3*) mutants were flood-inoculated at ∼10^6^ CFU/mL on *A. chinensis* var. *chinensis* ‘Hort16A’, and photographs of symptom development in pottles and representative leaves taken at 50 days post-inoculation.

**Supplementary Figure S6.**
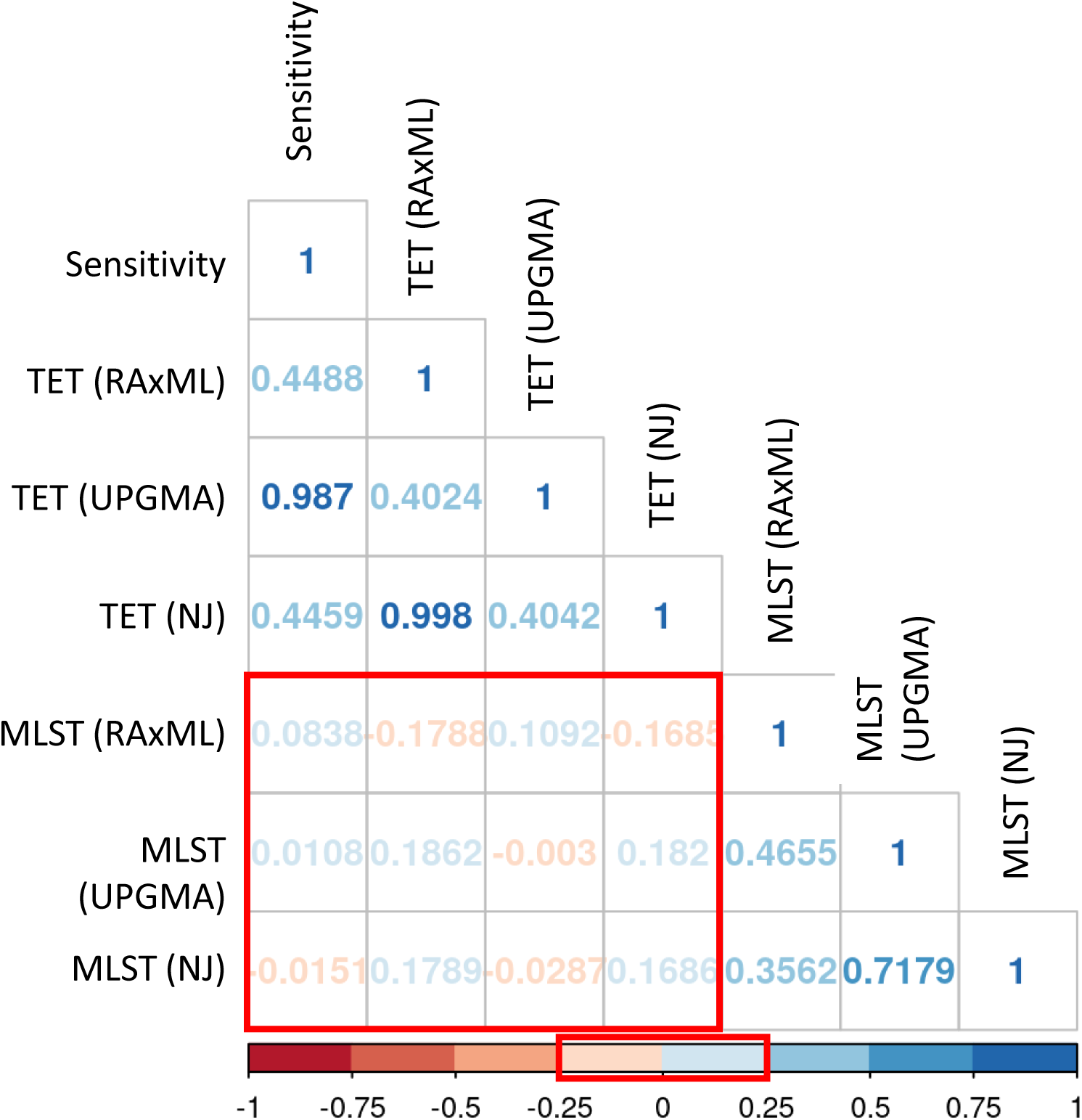
Cophenetic entanglement coefficients for phylogenetic trees for the TET operon and MLST analysis generated using different tree-building methods (UPGMA, Neighbor joining (NJ), or RAxML) compared to each other and to the R-syringacin (tailocin) sensitivity tree for data from Baltrus *et al.* (2019). The cophenetic correlation matrix between the trees have values ranging between -1 to 1, with near 0 values indicating that the two trees are not statistically similar. Values between -0.25 and 0.25 (indicated by the red box on the scale) are poorly correlated and outlined in the table and associated scale. UPGMA was selected over RAxML and NJ for tree-building in Fig. 6 due to the nature of comparing the sensitivity phenotype with genetic similarity treated as a ‘phenotype’ (thus creating phenetic trees rather than phylogenetic trees). This allowed for pairwise comparisons instead of inferring an evolutionary relationship that NJ and RAxML trees are better at conveying.

**Supplementary Table 1.** Metadata for isolates used in this study

| *P. syringae* isolate | Abbreviation | Strain | Host | Country of isolation | Date | GenBank accession |
| --- | --- | --- | --- | --- | --- | --- |
| *P. marginalis* pv. *marginalis* | *Pmar* | ICMP 3553PT | *Cichorium intybus* | New Zealand | 1949 | RBPW01 |
| *P. syringae* | *Ps* | CC1417 | Epilithon | USA |  | AVEO02 |
| *P. syringae* | *Ps* | CC1524 | Stream water | France |  | AVEK02 |
| *P. syringae* | *Ps* | CC1583 | Epilithon | France |  | AVEF02 |
| *P. syringae* | *Ps* | G33C | *Actinidia* sp. | New Zealand |  |  |
| *P. syringae* | *Ps* | ICMP 11296 | *Actinidia deliciosa* | New Zealand | 1991 | RBRK01 |
| *P. syringae* | *Ps* | ICMP 19498 | *Actinidia chinensis* | New Zealand | 2010 | LKCH01 |
| *P. syringae* | *Ps* | ICMP 3272 | *Actinidia deliciosa* | New Zealand | 1971 | LKEK01,<br>RBQZ01 |
| *P. syringae* | *Ps* | UB246 | Environmental | France |  | AVEQ01 |
| *P. syringae* | *Ps* | CC1416 | Epilithon | USA |  | AVEP02 |
| *P. syringae* | *Ps* | CC1458 | 02 | USA |  | AVEN02 |
| *P. syringae* | *Ps* | CC1466 | *Dodecatheon pulchellum* | USA |  | AVEM02 |
| *P. syringae* | *Ps* | CC1513 | *Pritzelago alpina* | France |  | AVEL02 |
| *P. syringae* | *Ps* | CC1544 | Lake water | France |  | AVEI02 |
| *P. syringae* | *Ps* | CC1557 | Environmental (snow) | France | ND | CP007014-5 |
| *P. syringae* | *Ps* | CC1559 | Environmental (snow) | France |  | AVEG02 |
| *P. syringae* | *Ps* | CC1630 | *Onobrychis viciifolia* | USA |  | AVED02 |
| *P. syringae* | *Ps* | CC440 | Cantaloupe | France |  | AVEC02 |
| *P. syringae* | *Ps* | CC457 | Cantaloupe | France |  | AVEB02 |
| *P. syringae* | *Ps* | CC94 | Cantaloupe | France |  | AVEA02 |
| *P. syringae* | *Ps* | ICMP 11168 | *Actinidia deliciosa* | New Zealand | 1991 | LKGV01 |
| *P. syringae* | *Ps* | ICMP 11292 | *Actinidia deliciosa* | New Zealand | 1991 | LKGU01 |
| *P. syringae* | *Ps* | ICMP 11293 | *Actinidia deliciosa* | New Zealand | 1991 | LKEP01 |
| *P. syringae* | *Ps* | ICMP 13102 | *Actinidia deliciosa* | France | 1985 | LKEO01 |
| *P. syringae* | *Ps* | ICMP 19198 | *Actinidia* sp. | New Zealand | 2011 | RBRZ01 |
| *P. syringae* | *Ps* | ICMP 19499 | *Actinidia* sp. | New Zealand | 2010 | LKCI01 |
| *P. syringae* | *Ps* | ICMP 3690 | *Prunus persica* | United Kingdom |  | LKBV01 |
| *P. syringae* | *Ps* | ICMP 4122 | *Prunus armeniaca* | New Zealand | 1974 | LKBX01 |
| *P. syringae* | *Ps* | TLP2 | Potato | USA |  | JGI<br>2507262033 |
| *P. syringae* | *Ps* | UB303 | Lake water | France |  | AVDZ |
| *P. syringae* | *Ps* | USA007 | Stream water | USA |  | AVDY02 |
| *P. syringae* | *Ps* | USA011 | Stream water | USA |  | CP045799-01 |
| *P. syringae* | *Ps* | CC1543 | Lake water | France |  | AVEJ02 |
| *P. syringae* pv. *actinidiae* BV1 | *Psa* BV1 | ICMP 9853 | *Actinidia deliciosa* | Japan | 1984 | CP018202-4 |
| *P. syringae* pv. *actinidiae* BV2 | *Psa* BV2 | ICMP 19071 | *Actinidia chinensis* | Korea | 1997 | AOJS01,<br>RBSG00 |
| *P. syringae* pv. *actinidiae* BV3 | *Psa* BV3 | ICMP 18884 | *Actinidia deliciosa* | New Zealand | 2010 | CP011972-3 |
| *P. syringae* pv. *actinidiae* BV5 | *Psa* BV5 | MAFF 212063 | *Actinidia chinensis* | Japan | 2012 | CP024712-4 |
| *P. syringae* pv. *actinidiae* BV6 | *Psa* BV6 | MAFF 212141 | *Actinidia deliciosa* | Japan | 2015 | MSBX01 |
| *P. syringae* pv. *actinidifoliorum* | *Pfm* L1 | ICMP 18802 | *Actinidia chinensis* | New Zealand | 2010 | MUKM01 |
| *P. syringae* pv. *actinidifoliorum* | *Pfm* L1 | ICMP 18803 | *Actinidia chinensis* | New Zealand | 2010 | AOKK01 |
| *P. syringae* pv. *actinidifoliorum* | *Pfm* L3 | ICMP 18807 | *Actinidia deliciosa* | New Zealand | 2010 | ANJL01,<br>AOKG01 |
| *P. syringae* pv. *actinidifoliorum* | *Pfm* | ICMP 19497 | *Actinidia chinensis* | New Zealand | ND | LKBQ01 |
| *P. syringae* pv. *actinidifoliorum* | *Pfm* | ICMP 19486 | *Actinidia chinensis* | Australia | 1990 |  |
| *P. syringae* pv. *actinidifoliorum* | *Pfm* L4 | CFBP 8039 | *Actinidia deliciosa* | France |  | LJJM01 |
| *P. syringae* pv. *actinidifoliorum* | *Pfm* L2 | CFBP 8043 | *Actinidia deliciosa* | France |  | LJFM01 |
| *P. syringae* pv. *actinidifoliorum* | *Pfm* L4 | CFBP 8160 | *Actinidia deliciosa* | France |  | LJJL01 |
| *P. syringae* pv. *actinidifoliorum* | *Pfm* L1 | CFBP 8161 | *Actinidia deliciosa* | France |  | LJFL01 |
| *P. syringae* pv. *actinidifoliorum* | *Pfm* L1 | CFBP 8180 | *Actinidia deliciosa* | France |  | LJFN01 |
| *P. syringae* pv. *aesculi* | *Pae* | 0893_23 | Horse chestnut | United States |  | AEAD01 |
| *P. syringae* pv. *amylgdali* | *Pam* | ICMP 3918 | *Prunus dulcis* | Greece | 1967 | LJPQ01 |
| *P. syringae* pv. *antirrhini* | *Pat* | ICMP 4959 | *Antirrhinum majus* | United Kingdom | 1965 | RBQK01 |
| *P. syringae* pv. *aptata* | *Ptt* | G733 | *Oryza sativa* | ND | 1976 | RBOI01 |
| *P. syringae* pv. *aptata* | *Ptt* | DSM50252 |  |  |  | AEAN01 |
| *P. syringae* pv. *atrofaciens* | *Paf* | DSM50255 | Wheat |  |  | AWUI01 |
| *P. syringae* pv. *avellanae* | *Pav* | BPIC 631 | *Corylus avellana* | Greece | 1976 | ATDK01 |
| *P. syringae* pv. *avellanae* | *Pav* | ISPave037 | *Corylus avellana* |  |  | AKCK01 |
| *P. syringae* pv. *berberidis* | *Pbe* | ICMP 13291 | *Berberidis* sp. | New Zealand | 1995 | RBQP01 |
| *P. syringae* pv. *cannibina* | *Pcb* | ICMP 2823 | *Cannabis sativa* | Hungary | 1957 | LJPX01,<br>FNKU01 |
| *P. syringae* pv. *cerasicola* | *Pce* | ICMP 13929 | *Prunus x yedoensis* | Japan | 1996 | LKCB01 |
| *P. syringae* pv. *cerasicola* | *Pce* | ICMP 17525 | *Prunus x yedoensis* | Japan |  | LKCC01,<br>RBTI01 |
| *P. syringae* pv. *cichorii* | *Pci* | ICMP 6917 | *Carthamus tinctorius* | New Zealand | 1980 | RBRY01 |
| *P. syringae* pv. *delphinii* | *Pde* | ICMP 529 | *Delphinium* sp. | New Zealand | 1957 | LJQH01 |
| *P. syringae* pv. *glycinea* | *Pgy* | ICMP 2236 | *Glycine max* | New Zealand |  | RBRO01 |
| *P. syringae* pv. *glycinea* | *Pgy* | 2411 | *Glycine max* | New Zealand | 1971 | QJTT01 |
| *P. syringae* pv. *glycinea* | *Pgy* | ICMP 2189 | *Glycine max* | New Zealand | 1968 | LJQL01 |
| *P. syringae* pv. *glycinea*<br>Race 4 | *Pgy* | A29-2 |  |  |  | ADWY01 |
| *P. syringae* pv. *lachrymans* | *Pla* | NM002 | Cucumber | China | 1984 | CP020351.1 |
| *P. syringae* pv. *lachrymans* | *Pla* | 98A-744 | *Cucumis sativus* |  |  | LCWT01 |
| *P. syringae* pv. *lachrymans* | *Pla* | MAFF 302278 |  |  |  | AEAM01 |
| *P. syringae* pv. *lachrymans* | *Pla* | MAFF370057 |  |  | 1979 | JGI<br>2505313044 |
| *P. syringae* pv. *lachrymans* | *Pla* | MAFF301315 | Cucumber |  |  | CP031225-8 |
| *P. syringae* pv. *lapsa* | *Plp* | ATCC 10859 | *Triticum aestivum* | ND | 2014 | CP013183 |
| *P. syringae* pv. *lapsa* | *Plp* | ICMP 3947 | *Zea sp.* | ND | ND | LJQQ01 |
| *P. syringae* pv. *maculicola* | *Pma* | 90-32 | *Brassica oleracea* | United States | 1990 | LGLH01 |
| *P. syringae* pv. *maculicola* | *Pma* | ICMP 11281 | *Brassica rapa* | China | ND | RBUQ01 |
| *P. syringae* pv. *mori* | *Pmo* | ICMP 4468 | *Morus alba* | New Zealand | 1957 | RBRW01 |
| *P. syringae* pv. *mori* | *Pmo* | MAFF 301020 | Mulberry | Japan | 1966 | AEAG01 |
| *P. syringae* pv.<br>*morsprunorum* | *Pmp* | NZIPFRPS-6 |  | New Zealand | 2010 | LKCD01 |
| *P. syringae* pv.<br>*morsprunorum* | *Pmp* | MAFF302280 |  |  |  | AEAE01 |
| *P. syringae* pv. *oryzae* | *Por* | ICMP 9093 | *Oryza sativa* | Japan | 1983 | RBSZ01 |
| *P. syringae* pv. *oryzae* | *Por* | 1_6 | *Oryza sativa* |  | 1991 | ABZR01 |
| *P. syringae* pv. *persicae* | *Ppe* | ICMP 3706 | *Prunus cerasifera* | New Zealand | 1966 | RBQE01 |
| *P. syringae* pv.<br>*phaseolicola* | *Pph* | 1448A | *Phaseolus vulgaris* | Ethiopia | 1985 | CP000058-60 |
| *P. syringae* pv.<br>*phaseolicola* | *Pph* | 1644 | *Vigna radiata* |  |  | AGAS01 |
| *P. syringae* pv. *pisi* | *Ppi* | 1704B |  |  |  | AEAI0 |
| *P. syringae* pv. *pisi* | *Ppi* | PP1 | *Pisum sativum* | Japan | 1978 | CP034078-81 |
| *P. syringae* pv. *savastanoi* | *Psv* | NCPPB 3335 | *Olea europaea* | France | ND | CP008742.1 |
| *P. syringae* pv. *syringae* | *Psy* | B48 | *Prunus persica* | United States |  | LGKT01 |
| *P. syringae* pv. *syringae* | *Psy* | 5264 | *Prunus avium* | United Kingdom | 2017 | NBAQ01 |
| *P. syringae* pv. *syringae* | *Psy* | B728a | *Phaseolus vulgaris* | United States | 1987 | CP000075.1 |
| *P. syringae* pv. *syringae* | *Psy* | PD 2766 | *Actinidia* sp. | United States |  | LKEM01 |
| *P. syringae* pv. *syringae* | *Psy* | PD 2774 | *Actinidia* sp. | United States |  | LKEL01 |
| *P. syringae* pv. *syringae* | *Psy* | B301D | *Pyrus communis* | United Kingdom | 1959 | CP005969 |
| *P. syringae* pv. *tabaci* | *Pta* | 6605 | Tobacco | Japan |  | AJXI01 |
| *P. syringae* pv. *tabaci* | *Pta* | ICMP 2835 | *Nicotiana tabacum* | Hungary | 1959 | LJRL01 |
| *P. syringae* pv. *tagetis* | *Ptg* | ICMP 4092 | *Tagetes erecta* | United kingdom | 1970 | RBQC01 |
| *P. syringae* pv. *tagetis* | *Ptg* | ICMP 6369 | *Tagetes erecta* | Australia | 1976 | RBVF01 |
| *P. syringae* pv. *thea* | *Pth* | NCPPB 2598 | *Thea sinensis* | Japan | 1974 | AGNN01 |
| *P. syringae* pv. *tomato* | *Pto* | DC3000 | *Solanum lycopersicum* | United Kingdom | 1960 | AE016853-5 |
| *P. syringae* pv. *tomato* | *Pto* | ICMP 2844 | *Solanum lycopersicum* | United Kingdom | 1960 | LJRN01 |
| *P. syringae* pv. *tomato* | *Pto* | T1 |  | Canada | ND | ABSM01 |
| *P. syringae* pv. *tremae* | *Ptr* | CC1629 | Oats | USA |  | AVEE02 |
| *P. viridiflava* | *Pvir* | CC1582 | Epilithon | France |  | AVDW01 |
| *P. viridiflava* | *Pvir* | ICMP 2848 | *Phaseolus vulgaris* | Switzerland | 1927 | LJRS01,<br>LKEH01 |
| *P. viridiflava* | *Pvir* | ICMP 8820 | *Prunus persica* | USA | 1984 | LKCA01 |
| *P. viridiflava* | *Pvir* | TA043 | *Primula officinalis* | France | ND | AVDV01 |
| *P. viridiflava* | *Pvir* | CDRTc14 | *Lepidium draba* | Austria | 2013 | MBPF01 |
| *P. viridiflava* | *Pvir* | ICMP 11289 | *Actinidia deliciosa* | New Zealand | 1991 | LKGX01 |
| *Pseudomonas* spp. | *P. spp.* | ICMP 10191 | *Actinidia* sp. | China | 1981 | LKGW01 |

**Supplementary Table 2.**
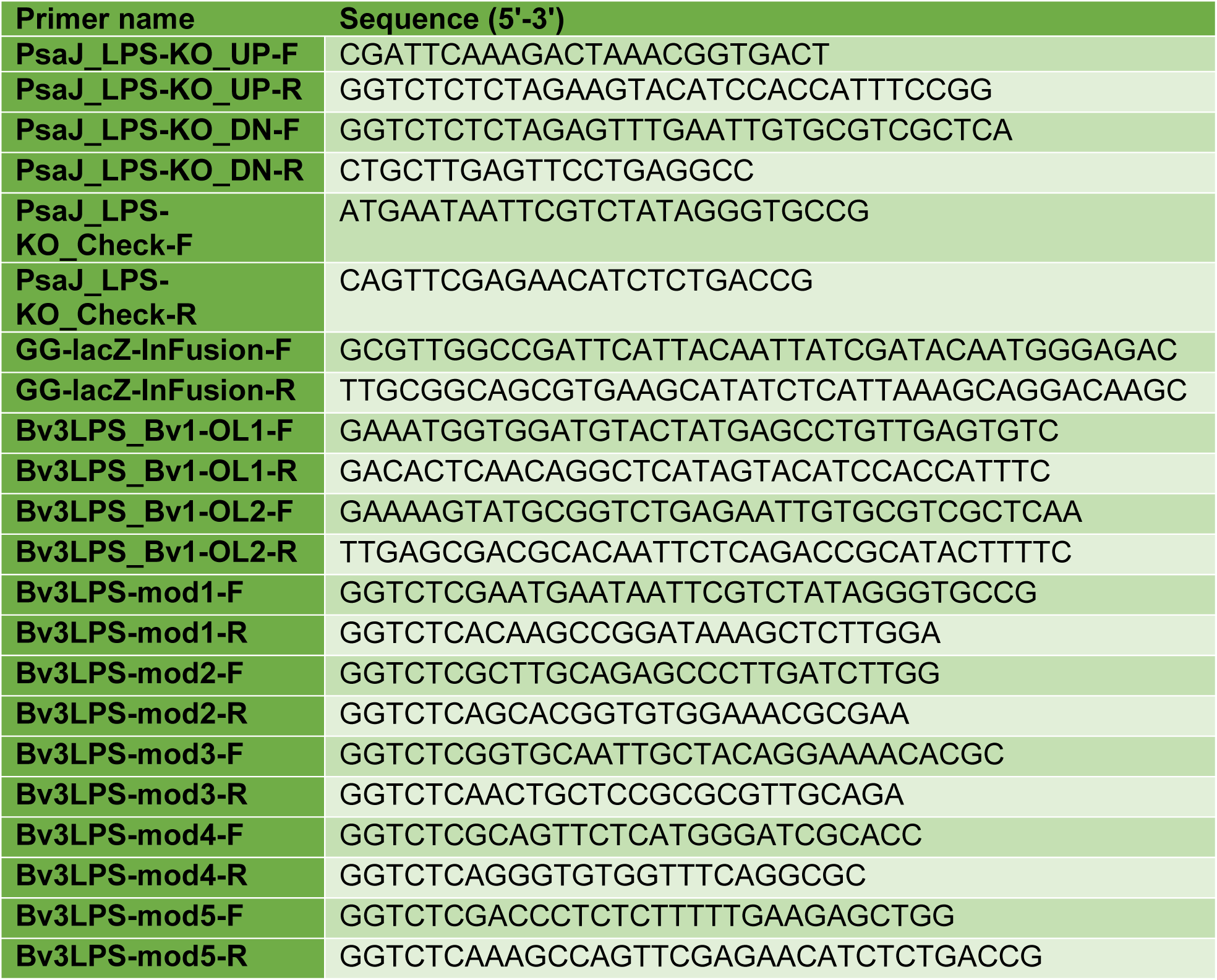
The sequence of primers used in this study. All primers were synthesized by Macrogen, South Korea.

